# Analysis of neuronal Ca^2+^ handling properties by combining perforated patch clamp recordings and the added buffer approach

**DOI:** 10.1101/2020.07.10.164202

**Authors:** Simon Hess, Christophe Pouzat, Lars Paeger, Andreas Pippow, Peter Kloppenburg

## Abstract

Ca^2+^ functions as an important intracellular signal for a wide range of cellular processes. These processes are selectively activated by controlled spatiotemporal dynamics of the free cytosolic Ca^2+^. Intracellular Ca^2+^ dynamics are regulated by numerous cellular parameters. Here, we established a new way to determine neuronal Ca^2+^ handling properties by combining the ‘added buffer’ approach (Neher and Augustine, 1992) with perforated patch-clamp recordings (Horn and Marty, 1988). Since the added buffer approach typically employs the standard whole-cell configuration for concentration-controlled Ca^2+^ indicator loading, it only allows for the reliable estimation of the immobile fraction of intracellular Ca^2+^ buffers. Furthermore, crucial components of intracellular signaling pathways are being washed out during prolonged whole-cell recordings, leading to cellular deterioration. By combining the added buffer approach with perforated patch-clamp recordings, these issues are circumvented, allowing the precise quantification of the cellular Ca^2+^ handling properties, including immobile as well as mobile Ca^2+^ buffers.

## Introduction

This study combined perforated patch clamp recordings with ratiometric Ca^2+^ imaging and the ‘added buffer approach’ to achieve significant improvements in the quantitative analysis of neuronal Ca^2+^ handling. In neurons, Ca^2+^ serves as a ubiquitous signal in a wide range of cellular processes, including the synaptic release of neurotransmitters, membrane excitability, enzyme activation, and activity-dependent gene activation (Augustine et al., 2003; Berridge, 1998; Clapham, 2007; Gilabert, 2020). These processes are selectively regulated by controlled spatiotemporal dynamics of the free cytosolic Ca^2+^ (Berridge, 2006; Berridge et al., 2000). Intracellular Ca^2+^ dynamics are regulated by numerous cellular parameters (Ca^2+^ handling parameters) including Ca^2+^ influx through voltage- and ligand-gated Ca^2+^ channels, Ca^2+^ release from intracellular stores, the location of the Ca^2+^ source, Ca^2+^ buffering by mobile and immobile buffers, Ca^2+^ extrusion, locally changing diffusion coefficients as well as the geometry of the cell (Helmchen et al., 1996; Kloppenburg et al., 2000; Lin et al., 2017; Maravall et al., 2000; Neher and Augustine, 1992; Sabatini et al., 2002; Tank et al., 1995). Impairment of this finely tuned system can cause neuronal dysfunction and even neurodegeneration (Berridge, 2011; Chan et al., 2009). Neuronal Ca^2+^ handling properties have been, up to now, analyzed by combining electrophysiological recordings and Ca^2+^ imaging into what is referred to as the ‘added buffer’ approach (Neher and Augustine, 1992).

The added buffer approach is based on a single-compartment model with the following rationale. During the measurement of intracellular Ca^2+^ dynamics with Ca^2+^ chelator-based indicators, the amplitude and time course of the Ca^2+^ signals (free cytosolic Ca^2+^ concentration) are influenced by the concentration of the Ca^2+^ indicator (e.g., fura-2), since the indicator acts as an exogenous (added) Ca^2+^ buffer and competes with the endogenous buffer(s). Accordingly, increasing indicator concentrations reduce the amplitude and prolong the decay time constants of free intracellular Ca^2+^ fluctuations (Helmchen et al., 1996; Neher nd Augustine, 1992; Pippow et al., 2009). If the buffering capacity (Ca^2+^ binding ratio) of the added buffer is known, the decay time constant can be used to estimate the Ca^2+^ signal kinetics in the absence of exogenous buffer. The model of the original added buffer approach (Neher and Augustine, 1992) provides estimates of Ca^2+^ handling parameters such as the free cytosolic Ca^2+^ concentration ([Ca^2+^]_i_), the Ca^2+^ extrusion rate (γ), the Ca^2+^ binding ratio (κ_B_), and the endogenous decay time constant (τ_endo_).

The added buffer approach relies on information about the precise exogenous Ca^2+^ buffer concentration in the neuron. Typically, the Ca^2+^ indicator concentration can be determined at any time during the experiment from loading curves. The neurons are loaded *via* the patch pipette, whose solution contains a known Ca^2+^ indicator concentration. During loading, fluorescence is monitored continuously, and it is assumed that when the fluorescence reaches a plateau, the indicator concentrations in the cell and the pipette are equal. Loading is usually performed in the whole-cell patch clamp configuration since this recording configuration provides low access resistance, ensuring that the solution in the patch pipette can freely interchange with the cytoplasm. Thus, the whole-cell configuration is ideally suited to “control” the composition of the intracellular milieu, including the Ca^2+^ indicator (added Ca^2+^ buffer) concentration (what is in the patch pipette ends up being what is in the neuron). However, the free exchange of molecules between cytoplasm and patch pipette disturbs intracellular signaling by washing out components of the cytosolic signaling system, including, for example, mobile cytosolic Ca^2+^ buffers. This not only impairs neuronal function but also causes estimation errors of the Ca^2+^ handling properties (Akaike, 1994; Akaike and Harata, 1994; Delvendahl et al., 2015; Lindau and Fernandez, 1986; Matthews et al., 2013).

For pure electrophysiological recordings, this issue can be minimized or even ignored by using the perforated patch clamp configuration originally introduced by Horn and Marty (Horn and Marty, 1988). In this recording configuration, electrical access between the recording pipette and the intracellular space is mediated by perforating agents that form channels or pores in the membrane. The washout of cytosolic components and the disruption of intracellular signaling pathways are then drastically reduced compared to the whole-cell configuration (Korn and Horn, 1989). The original and most commonly used perforating agents have been the antibiotic polyenes nystatin and amphotericin B, and the antibiotic polypeptide gramicidin (Akaike, 1994; Akaike and Harata, 1994; Hess et al., 2013; Horn and Marty, 1988; Klöckener et al., 2011; Könner et al., 2011; Kyrozis and Reichling, 1995). While polyenes and the peptide exhibit differences in their pore-forming mechanisms and ion selectivities (De Kruijff and Demel, 1974; Myers and Haydon, 1972; Russell et al., 1977; Tajima et al., 1996), their pores share key common properties: they are permeable to small molecules with a molecular weight up to ~200 Da, including monovalent ions (Kruijff et al., 1974; Kyrozis and Reichling, 1995; Urry, 1971), but they are neither permeable to divalent ions like Ca^2+^ nor to Ca^2+^ indicators like fura-2 (whose molecular weight is >600 Da). While the combination of perforated patch clamp recordings and the added buffer approach would be ideal, it cannot be implemented with the previous standard small pore-forming perforating agents.

To alleviate the “small pore size” problem, we tried β-escin as the perforating agent. β-escin is a saponin (Sirtori, 2001) and is structurally not related to the previous perforating agents. The action of saponins on biological membranes is concentration-dependent and complex. Most models of its action propose that membrane permeabilization is mediated by cholesterol complexation (formation of saponin-cholesterol complexes), which disturbs the normal membrane structure or, at high concentrations, removes the cholesterol from the membrane (Bangham et al., 1962; Böttger and Melzig, 2013). Since the pore size is generally larger and accordingly less selective compared to the ones of nystatin, amphotericin B and gramicidin, molecules of higher molecular weight (e.g., fluo-3, MW 770 Da) can pass through the cell membrane (Fan and Palade, 1998; Orta et al., 2012). Nevertheless, β-escin permeabilized membrane still constitutes an effective diffusion barrier to larger molecules like Ca^2+^ binding proteins (Fan and Palade, 1998; Sarantopoulos et al., 2004). In previous studies, β-escin has been successfully used to permeabilize smooth muscle cells (Kobayashi et al., 1989) and β-escin based perforated patch clamp recordings have been performed on cardiac myocytes (Fan and Palade, 1998) and different neuron types (Inoue et al., 2014; Sarantopoulos et al., 2004).

Here, we combined β-escin based perforated patch clamp recordings with the added buffer approach to measure intracellular Ca^2+^ handling properties. A thorough quantitative analysis of the data showed that this approach leads to a significantly improved quantification of the Ca^2+^ handling properties of individual neurons from brain slices of adult mice. The core experiments were performed on *substantia nigra* (SN) dopaminergic (DA) neurons from brain slices of adult mice. SN DA neurons are endogenous pacemakers and rely very heavily on intact, precisely orchestrated intracellular pathways, especially cytosolic Ca^2+^ signaling. Additionally, we performed a set of immunohistochemical experiments in cerebellar Purkinje cells, a cell type which is known for containing very high concentrations of calbindin D-28k, a mobile Ca^2+^ binding protein (Abe et al., 1992).

## Results

The goal of this study was to implement the ‘added buffer approach’, a powerful method to determine neuronal Ca^2+^ handling properties, in combination with perforated patch clamp recordings, which allow for electrophysiological recordings without compromising intracellular signaling. Since the typical perforating agents, nystatin, amphotericin B, and gramicidin, do not allow controlled loading of Ca^2+^ indicators, we explored the suitability of β-escin for this purpose. This study was performed in brain slices of mice, mainly on DA SN neurons. These neurons are particularly well suited to test this experimental approach since they are endogenous pacemakers (Fig. 1) that strongly rely on intact, precisely controlled Ca^2+^ signaling (Bean, 2007; Lacey et al., 1989; Richards et al., 1997).

**Figure 1.**
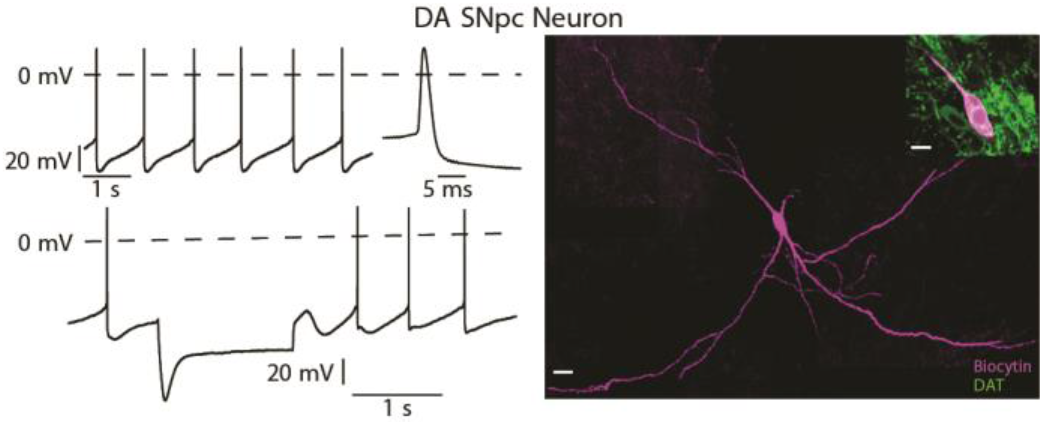
Basic identifying electrophysiological and morphological properties of dopaminergic substantia nigra neurons. The neurons were recorded in current clamp (*left*) and labeled with biocytin-streptavidin (magenta) *via* the patch pipette (*right*). The upper trace shows the regular pacemaker activity and long spike duration. On the lower trace, a hyperpolarizing current injection to −120 mV induced a clear ‘sag potential’, which is mediated by *I*_h_ and typical for this neuron type. Immunohistochemistry revealed dopamine transporter immunoreactivity (green), which a marker for dopaminergic neurons in the substantia nigra (*right, inset*). Scale bars: 20 μm, inset: 10 μm.

The first part of the study shows that β-escin enables long-lasting, high quality perforated, patch clamp recordings with minimal rundown. The quality of these recordings is comparable to amphotericin B perforated recordings. In the second part, we show that β-escin perforated patch clamp recordings allow controlled Ca^2+^ indicator loading and can significantly improve the added buffer approach to analyze the Ca^2+^ handling properties of individual neurons. We achieved controlled fura-2 loading with a minimized impact on the integrity of the cells’ cytoplasmic components. A thorough quantitative analysis of the data shows that this approach significantly improves the quantification of Ca^2+^ handling properties in single neurons of adult animals compared to conventional whole-cell recordings. The versatility of this new approach is complemented by immunohistochemical stainings of cerebellar Purkinje cells, which possess very high concentrations of the mobile Ca^2+^-binding protein calbindin D-28k. Here we could show that calbindin D-28k is retained during β-escin perforated patch clamp recordings, which is in stark contrast to results obtained during whole-cell patch clamp recordings.

### β-escin as perforating agent for perforated patch clamp recordings

After GΩ-seal formation, the perforation of the cell membrane was monitored by continuously measuring the membrane potential and the amplitude of the action potentials. Initially, ‘on-cell’ action potentials were observed in the cell-attached configuration (Fig. 2A). Diffusion of β-escin into the membrane gradually changed the recording to the perforated-patch configuration, decreasing the series resistance and increasing the action potential amplitude (Fig. 2A from left to right). The perforation process was considered fully established when action potential amplitude and series resistance (*R*_S_) had stabilized. Under our experimental conditions, we observed that indicator loading started within ~5 min of seal formation, and perforation was fully established after ~20 min (Fig. 2B).

**Figure 2.**
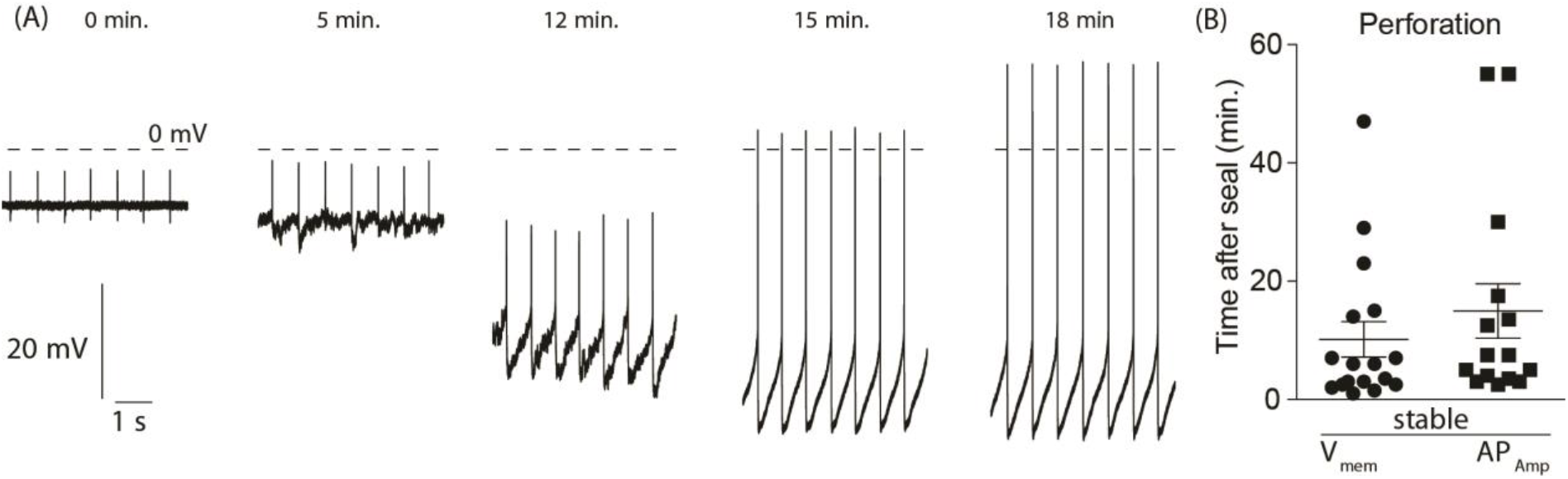
Perforated patch recordings of dopaminergic substantia nigra neurons using β-escin as a perforating agent. (A) Perforation process. Current clamp recording during the β-escin mediated membrane perforation showing the transition from ‘on-cell’ to the ‘perforated-patch’ configuration. The time (in min) after the seal was established, is given above each trace. (B) “Duration” of the perforation process: the time required for the baseline of the membrane potential (*V*_mem_) and the action potentials amplitude (AP_Amp_) to stabilize.

### Comparison with other patch clamp configurations

To compare the β-escin mediated recordings with other configurations of patch clamp recordings, we analyzed whether and how important intrinsic neurophysiological parameters such as spontaneous or evoked action potential firing, membrane potential, and resting Ca^2+^ levels ([Ca^2+^]_i_) are affected by the different recording configuration (whole-cell recordings, and amphotericin B- and β-escin perforated patch recordings). After establishing recordings with stable access (as judged by the series resistance), spontaneous and evoked action potential firing was monitored for at least 30 min. During whole-cell recordings, we observed a dramatic rundown of these parameters. As expected, they did not change during amphotericin B perforated-patch recordings. β-escin perforated-patch recordings gave results similar to amphotericin B mediated recordings, and only a small decline of the spontaneous firing was observed (Fig. 3A-D).

**Figure 3.**
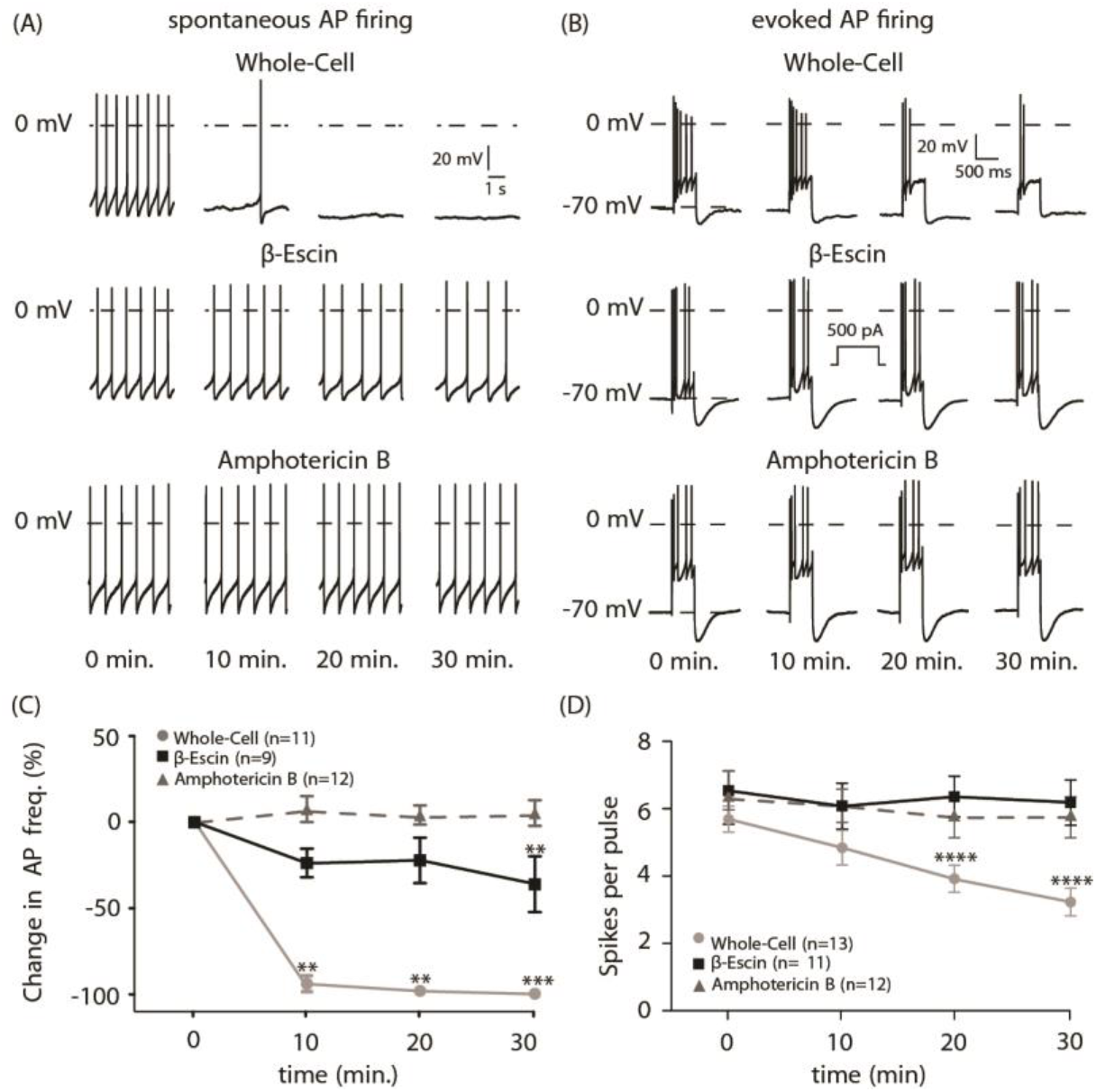
β-escin mediated perforated patch clamp recordings allow stable recordings. (A,B) spontaneous activity (A) and depolarization-induced spike trains (B) recorded from DA SN neurons in the whole-cell and perforated patch clamp (β-escin- or amphotericin mediated) configuration at different times after reaching “stable” recording conditions. (C,D) Stability of electrophysiological parameters in the different recording configurations. (C) Spontaneous activity. Friedman-test with post-hoc Dunn’s multiple comparison test. (D) The number of action potentials per current pulse (holding potential −70 mV; current pulse, 500 pA for 500 ms). Note that the large hyperpolarization following the current injection disappeared during the whole-cell recording. Repeated measures one-way ANOVA with Dunnett’s multiple comparison test. **: p≤0.01, ***: p≤0.001, ****: p≤0.0001.

More specifically, during whole-cell recordings, more than 80% of the cells stopped spontaneous firing within 10 min after break-in (Fig. 3A, C). This loss of activity was accompanied by a ~4 fold increase in the resting [Ca^2+^]_i_ levels after 30 min of recording time (Fig. 4). This substantial rise in [Ca^2+^]_i_ was a clear and direct indicator for the deteriorating physiological state of the neurons (Kuchibhotla et al., 2008; Mattson, 2007) and showed directly that the Ca^2+^ handling properties change during whole-cell recordings. In contrast, [Ca^2+^]_i_ remained stable during β-escin perforated patch recordings over the same time (Fig 4). Taken together, β-escin perforated patch clamp recordings significantly outperformed whole-cell recordings, and β-escin perforated patch clamp recordings were of similar quality to amphotericin B mediated recordings. Overall, the data are in line with many earlier studies (e.g. (Horn and Marty, 1988; Husch et al., 2011; Lindau and Fernandez, 1986)) which clearly show the impact of whole-cell recordings on the cytosolic integrity and to what extend perforated patch clamp can minimize this impact (rundown), and thus are suitable to analyze ‘true’ intrinsic neurophysiological parameters over long periods of time.

**Figure 4.**
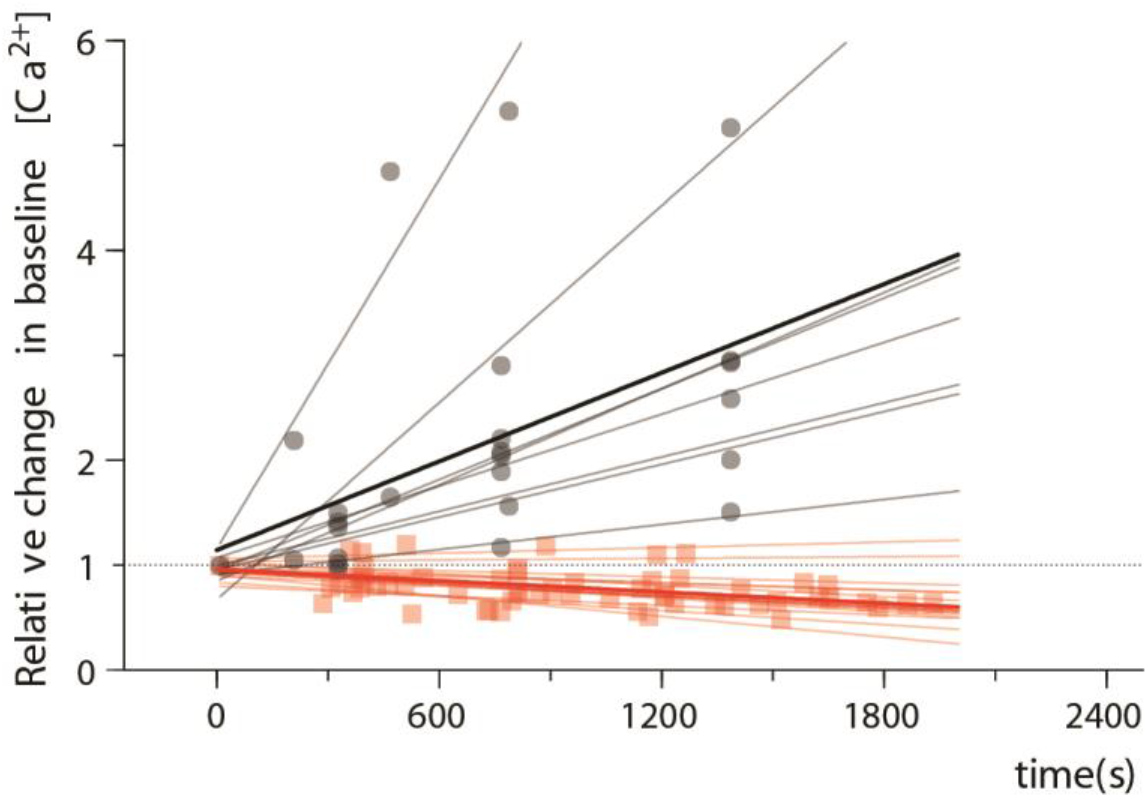
Intracellular Ca^2+^ concentrations remain stable during β-escin mediated perforated patch clamp recordings. Relative changes in intracellular Ca^2+^ concentration during the time course of β-escin perforated (pale red, red) and whole-cell patch clamp recordings (grey, black). Linear fits of all experiments (pale red and grey) are shown, with individual data points represented as squares. The bold red and black line are the linear fits of all β-escin perforated and whole-cell recordings. Whole-cell n=8; β-escin n=13. p<0.0001; F-test.

### Permeability of β-escin mediated pores

Next, we tested if the β-escin perforated membranes are permeable to fura-2 and to molecules with a MW in the range of Ca^2+^ buffering proteins, like calbindin D-28k. To clarify this question, tetramethylrhodamine-dextran (TRD-40kDa, MW 40 kDa) or fura-2 was added to the pipette solution, and the fluorescence in the cell body was monitored during the recording. During β-escin perforated-patch recordings, the TRD-40kDa remained in the recording pipette, while it diffused into the neuron during whole-cell recordings (Fig 5). Furthermore, immunohistochemical staining showed that the endogenous calbindin-D28k remains in the cell during β-escin perforated patch clamp recordings but is washed out during whole-cell recordings (Supplement Fig. 1). In contrast, the loading curves for fura-2 (Fig. 6A) confirmed that fura-2 could diffuse readily through the β-escin perforated membrane into the cell. These results suggested that β-escin is well suited to implement the added buffer approach.

**Figure 5.**
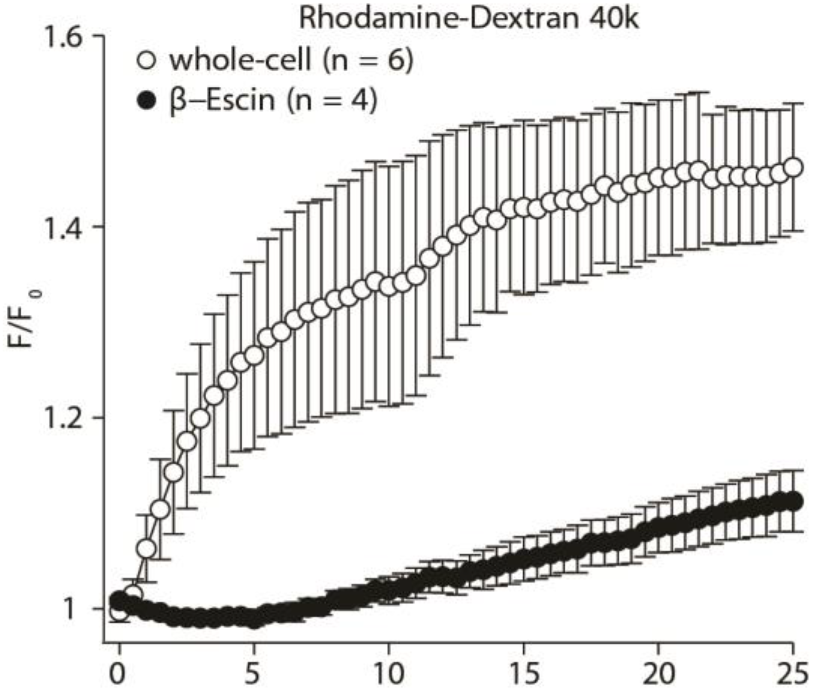
Large molecules diffuse very slowly through the perforated membrane during patch clamp recordings with β-escin. Time course of tetramethylrhodamine-dextran (40 kDa) fluorescence in DA SN neurons for whole-cell (○) and ß-escin perforated patch recordings (●). n-values are given in the figure. Values are mean ± SEM.

**Figure 6.**
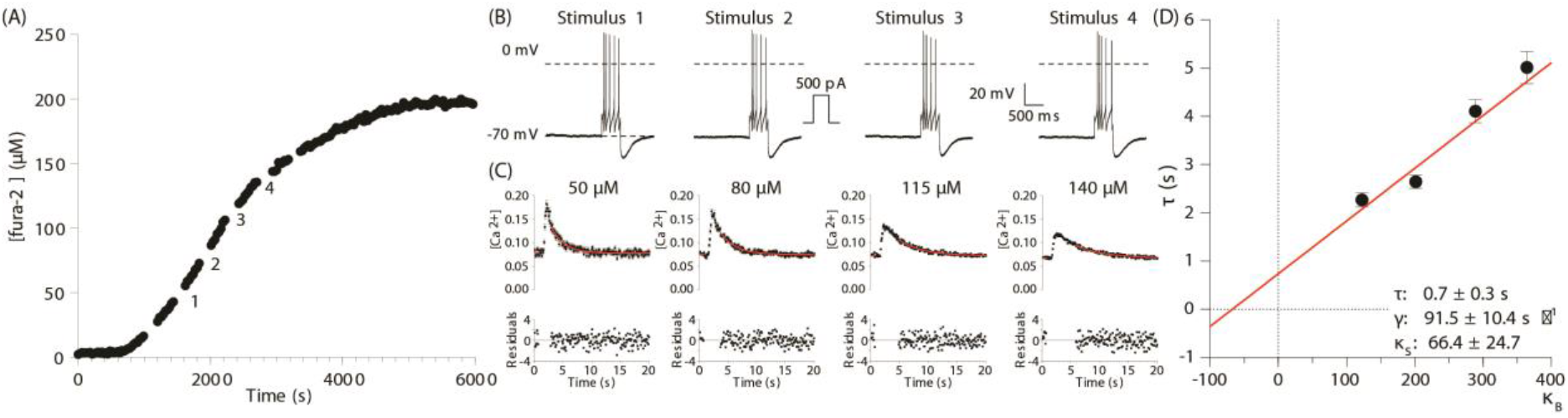
Outline of the Ca^2+^ handling properties analysis. Ca^2+^ handling was analyzed using a combination of patch clamp recordings, ratiometric Ca^2+^ imaging, and the ‘added buffer approach’. (A-D) demonstrate the principal steps of the analysis for a single DA SN neuron. The neuron was recorded and fura-2-loaded in the β-escin perforated patch clamp configuration. (A) Fura-2 loading curve. Neurons were loaded via the patch pipette with the ratiometric Ca^2+^ indicator fura-2, which also serves as the added Ca^2+^ buffer. Fura-2 fluorescence was acquired at 360 nm excitation (isosbestic point of fura-2) every 30 s and converted into fura-2 concentrations. (B) Spike trains that induce the voltage-activated Ca^2+^ influx shown in (C) at different times during loading, as indicated in (A). (C) Monoexponential fits of the individual Ca^2+^ transients following the four stimulations. The graphs demonstrate the effect of the increasing added Ca^2+^ buffer (fura-2) concentration on the amplitude, and decay kinetics of voltage evoked transients of free Ca^2+^: the amplitude decrease while the decay time increases. Notice the error bars on the individual data points. These error bars allow us to properly normalize the residuals (the difference between observed and fitted values divided by the standard deviation) shown below. If the mono-exponential model is correct and if its parameters have been properly estimated, these residuals should be (approximately) drawn for a Gaussian distribution with mean 0 and SD 1. (D) The decay time constants were plotted against the Ca^2+^ binding ratios of fura-2 (κ_B_). κ_B_ was calculated from the intracellular fura-2 concentration, the *K*_d_ of fura-2, and the resting concentration of free intracellular Ca^2+^. The solid red line is the linear fit to the data. An estimate of κ_S_ was obtained as the negative x-axis intercept. The Ca^2+^ extrusion rate is estimated from the slope of the fit and the endogenous decay time constant from the intercept with the y-axis. The resulting Ca^2+^ handling parameters of this example are: τ_endo_: 0.7 ± 0.3 s, γ: 91.5 ± 10.4, s^−1^, κ_s_: 66.4 ± 24.7.

### Ca^2+^ handling properties

Given the stable and reliable recording conditions provided by β-escin, we combined β-escin perforated patch clamp recordings with ratiometric fura-2 Ca^2+^ imaging to analyze Ca^2+^ handling properties. Using the added buffer approach, we determined the resting Ca^2+^ concentration ([Ca^2+^]_i_), the Ca^2+^ binding ratio κ_S,_ and the extrusion rate γ of SN DA neurons, which display Ca^2+^ dependent endogenous pacemaking properties (Liu et al., 2014; Neuhoff et al., 2002; Smits et al., 2013). Neurons were held at −70 mV to avoid spontaneous spiking. Clear and reproducible Ca^2+^ influx could be induced by trains of action potentials elicited by current pulses (500 pA, 500 ms) (Fig. 6B). The resulting intracellular Ca^2+^ signals were monitored by ratiometric Ca^2+^ imaging (Fig 6C), and the data quality was compared with data acquired with conventional whole-cell patch clamp recordings.

Fura-2 loading: The intracellular concentration of the exogenous buffer at any time during the experiment was determined from the loading curves of the calcium indicator (Fig. 6A). DA SN neurons were loaded *via* the patch pipette with solutions containing 100 – 200 μM fura-2. During dye loading, the fluorescence of the cell body was monitored in 30 s intervals using excitation at 360 nm (isosbestic point) to determine the dye concentration. After the β-escin perforated-patch configuration was fully established, fluorescence appeared in the cell bodies, and fluorescence intensity increased until it reached a stable plateau. The intracellular fura-2 concentration was estimated for different times during the loading curve, assuming that cells were fully loaded when the fluorescence reached a plateau and the fura-2 concentration in the cell and the pipette is equal at this point (for details see Methods).

Ca^2+^ handling properties: If the buffer capacity of the added buffer is known, the time constant of the transient decay (*τ*_transient_) can be used to estimate by extrapolation the Ca^2+^ signal to conditions, with only endogenous buffers present (−*κ*_B_ = 1 + *κ*_S_). The model used for this study (Eq. (14), (Neher and Augustine, 1992) assumes that the decay time constants *τ*_transient_ are a linear function of the Ca^2+^ binding ratios (*κ*_B_ and *κ*_S_). *κ*_S_ was determined from the negative x-axis intercept of a *τ*_transient_ over *κ*_S_ plot (Fig. 6D). The slope of the straight-line fit is the inverse of the extrusion rate (*γ*). The point of intersection of the linear fit with the y-axis denotes the endogenous decay time constant *τ*_endo_ (no exogenous Ca^2+^ buffer in the cell). This is demonstrated in Fig. 6 with an example of a DA SN neuron with following Ca^2+^ handling parameters: τ_endo_: 0.7 ± 0.3 s, γ: 91.5 ± 10.4, s^−1^, κ_s_: 66.4 ± 24.7.

### Comparison of data quality between whole-cell and β-escin perforated patch clamp recordings

The data quality of the Ca^2+^ imaging experiments used to determine the Ca^2+^ handling properties was markedly better for the β-escin perforated-patch configuration compared to whole-cell recordings. Obvious evidence of “low-quality” data were implausible results such as negative Ca^2+^ binding coefficients or Ca^2+^ transients that did not return to baseline as frequently observed by us and previous studies (Fierro and Llano, 1996; Pippow et al., 2009). For data sets that did not show such obvious flaws, an analysis on a finer scale confirmed the significantly higher data quality of the β-escin perforated records. For this purpose, parameters, which characterize the quality of data sets of individual neurons, were quantified. This included the goodness of fit for each Ca^2+^ transient (a mono-exponential fit) and the linear regression that was used to fit the *τ*_transient_ − *κ*_B_ relation.

First, we tested the goodness of the mono-exponential fit for each Ca^2+^ transient. Since our analysis method yields a meaningful standard error for the estimated [Ca^2+^]_i_ (see Methods), a fit obtained with a weighted nonlinear least-square produces standard errors for the estimated parameters (basal [Ca^2+^]_i_, Δ[Ca^2+^]_i_ and, most importantly here, *τ*_transient_) as well as residuals following a known distribution if the data do indeed follow the model; namely the residual sum of squares (RSS) should be the realization of a draw from a χ^2^ distribution with a known number of degrees of freedom, and the residuals should not be correlated. We decided to keep a transient for further analysis if both the p-values of the RSS and the auto-correlation at lag 1 were larger than 0.01 (giving a probability of false-negative of 0.02). Since the decay fits were started once the transient had lost 50% of the peak increase (see Materials and Methods), the number of data points used for each fit is not fixed, meaning that the number of degrees of freedom of the χ2 distribution associated with each RSS is also changing. A proxy of the RSS that is easy to compare from transient to transient is the RSS per degree of freedom (that should be close to 1 for a good fit) and is illustrated in Fig. 7C. This goodness of fit criterium was fulfilled by 75% (24/32) of the transients measured in the whole-cell configuration compared to 92% (67/73) of the transient measured in the perforated configuration (Fig. 7D; using an exact test, based on the hypergeometric distribution, the probability that these two samples arise from the same underlying distribution is found to be smaller than 0.02). We then used the data sets with at least three ‘good’ transients to perform the linear regression (*τ*_transient_ − *κ*_B_ relation) and found that 15/15 (100 %) data sets with perforated recordings lead to an acceptable regression but only 2/6 (33 %) of those with whole-cell recordings (Fig. 7G). Thus, while experiments in the whole-cell configuration can yield results comparable to β-escin experiments (Fig. 8A [*right panel*]; κ_s_: 19.9 ± 25.1, τ_endo_: 0.4 ± 0.5 s, γ: 55.2 ± 10.4 s^−1^), it is evident that in many cases the whole-cell results were implausible (Fig. 8A, *middle and left panel*). In these cases, the linear regression fit of the data intercepts the x-axis at positive value leading to a negative Ca^2+^ binding ratio κ_s_ and a negative endogenous decay time constant τ_endo_, respectively (see (Delvendahl et al., 2015))(κ_s_: −72.9 ± 14.3, τ_endo_: −17.9 ± 2.7 s, γ: 4.0 ± 0.5 s^−1^, *middle panel;* κ_s_: −3.1 ± 10.3, τ_endo_: −0.04 ± 0.2 s, γ: 58.0 ± 4.4 s^−1^, *left panel*). It is important to note that the implausibility of the results from individual cells can be masked if pooled data sets from multiple neurons are analyzed (Fig. 8C and D).

**Figure 7.**
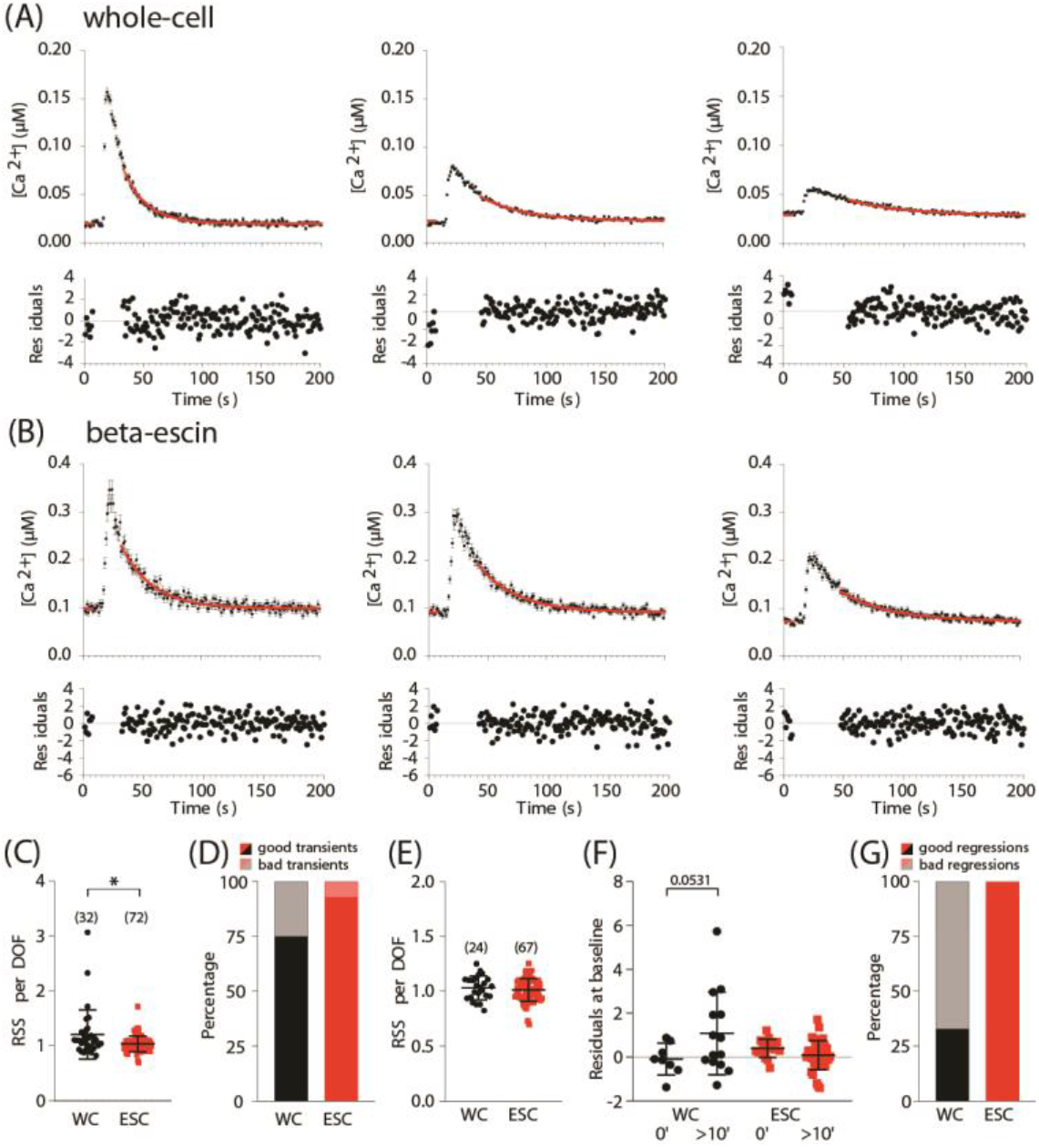
The goodness of fit comparison between Ca^2+^ transients recorded in the whole-cell and β-escin perforated patch clamp configuration. In both configurations, SN neurons were loaded with fura-2 via the recording pipette, as illustrated in figure 6. (A, B) Induced Ca^2+^ transients with increasing added Ca^2+^ buffer (fura-2) concentrations (from left to right). The monoexponential fit is shown in red with the residuals below. (C) Comparison of Residual Sum of Squares (RSS) per Degrees Of Freedom (DOF) between all recorded Ca^2+^ transients under both recording configurations; if everything goes well, a value close to 1 should be obtained. (D) Percentage of ‘good transients’ vs ‘bad transients’ (E) RSS per DOF of the ‘good transients’ shown in (D). (F) Comparisons of the residuals of the baseline fits at the beginning of the recording and more than 10 minutes after the beginning of the recording for both recording configurations. (G) Percentage of data sets with at least three ‘good transients’ that provided acceptable regressions.

**Figure 8.**
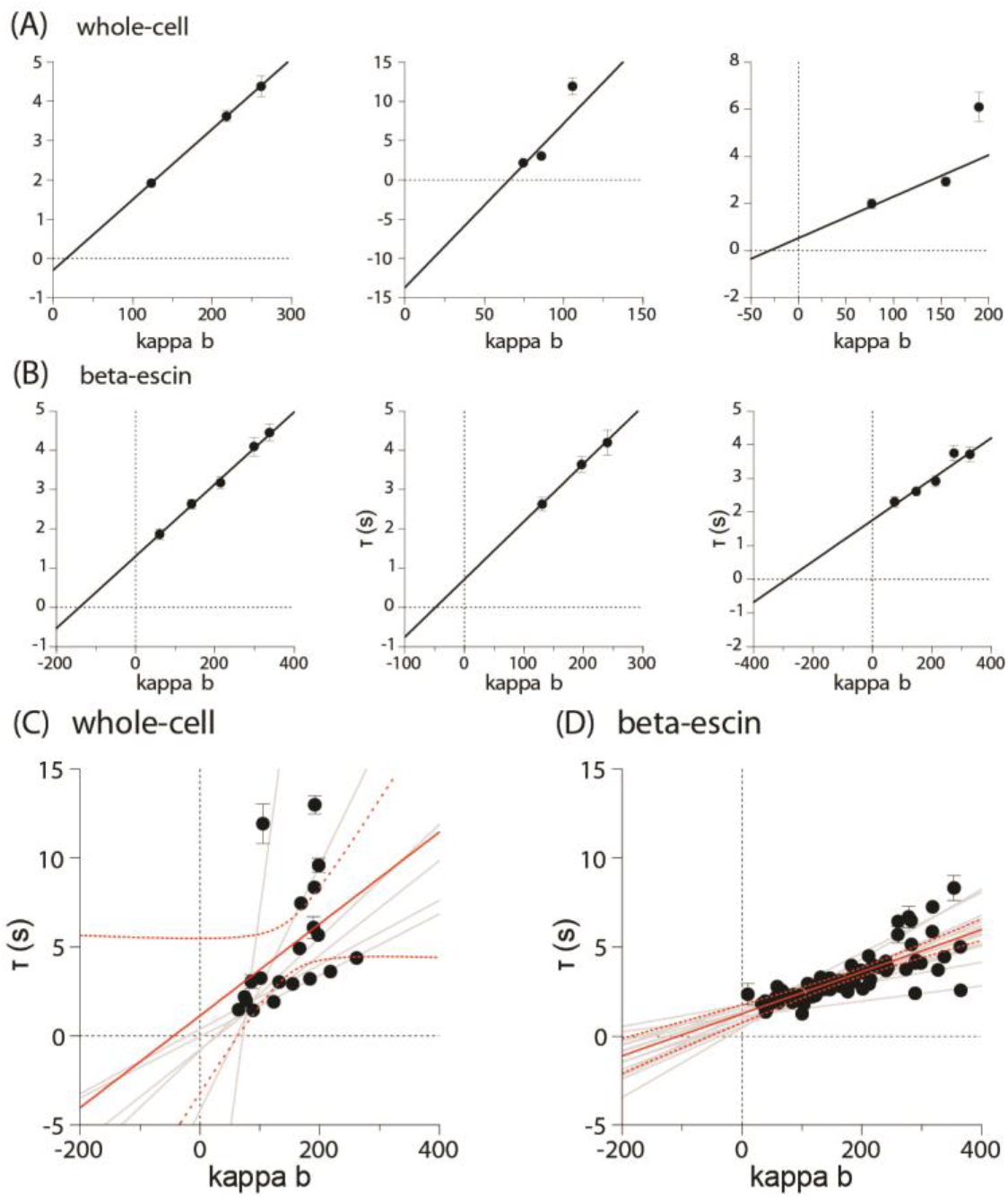
β-escin perforated patch clamp recordings provide systematically plausible Ca^2+^ handling properties. (A,B) τ_transient_ − κ_B_ relations of ‘added buffer’ experiments using the whole-cell (A) and β-escin perforated patch clamp configuration (B). (A) Data sets of three neurons obtained using the whole-cell configuration, demonstrating that this approach can provide plausible (right panel) and implausible (middle and left panel) results. (B) Three data sets that were obtained using the β-escin perforated configuration. Note the consistently ‘better fits’ and the higher plausibility of the x-axis intercept compared to (A). (C,D) τ_transient_ − κ_B_ relations of 6 SN neurons recorded in the whole-cell configuration (C) and 15 SN neurons recorded in the β-escin perforated patch clamp configuration (D). To demonstrate that the implausibility of individual data sets can be masked if pooled data sets of multiple neurons are analyzed, fitting was performed for the pooled data and for each neuron individually. For the pooled data, the best linear fits with 95% confidence bands are shown (solid and dashed red lines).

In summary, the overall quality of the data obtained in β-escin perforated patch clamp experiments is tremendously improved compared to experiments in the whole-cell configuration. This improvement results from two factors. The perforated configuration preserves the general integrity and functionality of the cytoplasmic pathways, including the Ca^2+^ handling machinery. In addition, it optimizes the measurements directly by preventing an uncontrolled change in mobile Ca^2+^ buffer during the recording. Ultimately, this resulted in a far more reliable and better-defined estimation of the Ca^2+^ handling properties of single neurons.

## Discussion

Using β-escin as a perforating agent, we showed that it is possible to combine longlasting perforated-patch recordings with controlled Ca^2+^ indicator loading and the ‘added buffer approach’ to analyze neuronal Ca^2+^ handling properties. We consider this a substantial progress, and of particular importance for the analysis of intracellular Ca^2+^ handling since the perforated-patch clamp configuration allows stable recordings with minimized impact on the intracellular signaling system, including mobile components of the Ca^2+^ handling machinery, i.e., mobile Ca^2+^ buffers. β-escin perforated recordings performed similarly well as amphotericin B, gramicidin, or nystatin perforated patch recordings, which are the ‘gold standards’ for stable long-term recordings (Horn and Marty, 1988; Husch et al., 2011; Rae et al., 1991). β-escin perforated patch recordings clearly outperformed conventional whole-cell recordings, which becomes especially apparent by monitoring the Ca^2+^ levels. In contrast to the whole-cell configuration, the resting Ca^2+^ concentration remained stable, indicating healthy recording conditions.

Typically, the ‘added buffer’ approach has been used in combination with conventional whole-cell patch clamp recordings, which allows the controlled loading of cells with Ca^2+^ indicator dyes (e.g., fura-2) via the patch pipette (Delvendahl et al., 2015; Lee et al., 2000a; Lin et al., 2017; Neher and Augustine, 1992). In this context, the whole-cell configuration has the obvious advantage that it allows for easy and reliable loading of the cell with Ca^2+^ indicators. However, this advantage is offset by the disadvantage that intracellular components are being washed out, leading to a gradual change of cellular properties over time. This causes not only general damaging effects on the physiology and Ca^2+^ handling machinery of the cell, but it also introduces a systematic error to the analysis, since in the whole-cell configuration endogenous mobile Ca^2+^ buffer is leaving the neuron at an unknown rate during loading of the ‘added’ buffer (here fura-2). Although this is not a new finding, since Neher and Augustine had already explicitly mentioned this matter in their original paper (Neher and Augustine, 1992), and although many studies have applied the added buffer approach in combination with whole-cell patch clamp recordings, this issue has not been addressed directly. In a strict sense, the added buffer approach in combination with the whole-cell configuration can only be used to analyze the immobile fraction of Ca^2+^ buffers and only if the measurements start *after* the mobile Ca^2+^ buffers have diffused out of the cell and do not interfere with the analysis. Accordingly, this approach can be explicitly used to analyze the immobile Ca^2+^ buffers as convincingly shown by Müller et al. (2005) and Matthews et al. (2013) in hippocampal neurons.

Employing β-escin as perforating agent addressed this issue and allowed to successfully combine perforated patch clamp recordings with controlled Ca^2+^ indicator loading, and thus the added buffer approach. While our and other’s whole-cell experiments (Delvendahl et al., 2015; Matthews et al., 2013; Müller et al., 2005) showed that molecules with a molecular weight comparable to Ca^2+^ binding proteins started being washed out immediately after establishing the whole-cell configuration, our permeability measurements showed that the β-escin perforated membranes retained TRD-40kDa in the recording pipette and calbindin D-28k in the neuron, indicating that the perforated membrane is impermeable for the relative large endogenous mobile Ca^2+^ buffers. Fura-2, on the other hand, with its low molecular weight, could easily permeate through the perforated membrane.

For practical purposes, it is important to note that the permeability of the perforated membrane is concentration-dependent. Thus, the β-escin concentration might have to be adjusted to different experimental conditions and cell types. Here, with β-escin concentrations of 10–15 μM, the perforated membrane was permeable to fura-2 but not to TRD-40kDa and calbindin D-28k. In permeabilized muscle cells, β-escin concentrations of 50 – 100 μM render cell membranes permeable to molecules ranging from 10 to 150 kDa MW (Iizuka et al., 1997, 1994; Konishi and Watanabe, 1995). In recordings of cardiac myocytes with 50 μM β-escin, the membrane was permeable for fluorescent markers with a MW up to 10 kDa after 20 – 25 min recording time (Fan and Palade, 1998).

Taken together, this study shows that the newly established approach is suitable to improve the quantitative analysis of intracellular Ca^2+^ handling parameters markedly. The combination of perforated patch clamp recordings, which preserve the integrity and functionality of the intracellular milieu, with the ‘added buffer approach’, is a powerful tool to analyze mobile and immobile components of the intracellular Ca^2+^ handling machinery.

## Materials and Methods

### Animals and brain slice preparation

All animal procedures were conducted in compliance with protocols that were approved by local government authorities (LANUV NRW, Recklinghausen, Germany). Mice were housed at 22 – 24 °C with a 12-hour light/12 hour dark cycle. Animals had access to water and chow *ad libitum*. Experiments were performed on brain slices from 9 – 11 weeks old male C57BL/6 mice. Preparation of brain slices, recordings from DA SN and Purkinje neurons, labeling, and identification of neurons was performed as previously described (Hess et al., 2013; Könner et al., 2011; Murru et al., 2019). The mice were lightly anesthetized with isoflurane (B506; AbbVie Deutschland GmbH & Co KG, Ludwigshafen, Germany) and subsequently decapitated. Coronal slices (250 – 300 μm) containing the SN or sagittal slices containing the cerebellum (300 μm) were cut with a vibration microtome (HM-650 V; Thermo Scientific, Walldorf, Germany) under cold (4°C), carbogenated (95% O_2_ and 5% CO_2_), glycerol-based modified artificial cerebrospinal fluid (GaCSF; (Ye et al., 2006)). GaCSF contained (in mM): 250 Glycerol, 2.5 KCl, 2 MgCl_2_, 2 CaCl_2_, 1.2 NaH_2_PO_4_, 10 HEPES, 21 NaHCO_3_, 5 Glucose, adjusted to pH 7.2 with NaOH. Brain slices were transferred into carbogenated artificial cerebrospinal fluid (aCSF). aCSF contained (in mM): 125 NaCl, 2.5 KCl, 2 MgCl_2_, 2 CaCl_2_, 1.2 NaH_2_PO_4_, 21 NaHCO_3_, 10 HEPES, 5 glucose, adjusted to pH 7.2 with NaOH. First, they were kept for 20 min in a 35°C ‘recovery bath’ and then stored at room temperature (~24°C) for at least 30 min prior to recording. During the recordings, the slices were continuously superfused with carbogenated aCSF. To block synaptic currents, aCSF contained 10^−4^ M PTX, 5 × 10^−5^ M D-AP_5_, and 10^−5^ M CNQX. All recordings were carried out at 24°C.

### Electrophysiological recordings

DA SN neurons and the cerebellar Purkinje cells were recorded using whole-cell and perforated patch clamp recordings. Neurons were visualized and imaged with a fixed stage upright microscope (BX51WI, Olympus, Hamburg, Germany) using a 20× water-immersion objective (XLUMPLFL, 0.95 numerical aperture, 2 mm working distance, Olympus) with a 4× magnification changer (U-TVAC, Olympus), infrared differential interference contrast optics (Dodt and Zieglgänsberger, 1990) and fluorescence optics. Neurons were pre-identified by the size and location of their somata. This pre-identification was verified in each case by a physiological characterization during the recording and *post hoc* by immunohistochemistry for cell type-specific markers. DA SN neurons were recorded in the ventral tier SN. They have a low frequency, regular firing patterns with broad action potentials, and generate a clear *I*_h_-dependent “sag”-potential upon hyperpolarization (Fig. 1) (Lacey et al., 1989; Richards et al., 1997; Ungless et al., 2001). Purkinje cells were identified by their distinct morphology and localization within the cerebellar cortex and by their high-frequency spontaneous firing and very narrow action potentials (~200 μs; Supplement Fig. 1A) (Bean, 2007; McKay and Turner, 2005, 2004)). During the experiments, biocytin (0.1%; B4261; Sigma) diffused through the broken or perforated membrane into the recorded cell. Post recording, the identity of DA SN neurons and cerebellar Purkinje cells was confirmed by their immunoreactivity for the dopamine transporter (Fig. 1) (Ciliax et al., 1995) or the calcium-binding protein calbindin D-28k (Supplement Fig. 1) (Abe et al., 1992; Fournet et al., 1986), respectively. Biocytin-streptavidin labeling combined with dopamine transporter or calbindin D-28k immunohistochemistry was performed as previously described (Hess et al., 2013; Murru et al., 2019).

### Whole-cell patch and perforated patch-clamp recordings

Electrodes with tip resistances between 3 and 5 MΩ were fashioned from borosilicate glass (0.86 mm inner diameter; 1.5 mm outer diameter; GB150-8P; Science Products, Hofheim, Germany) with a vertical pipette puller (PP-830; Narishige, London, UK). Recordings were performed with an EPC10 patch-clamp amplifier (HEKA, Lambrecht, Germany) controlled by the program PatchMaster (version 2.32; HEKA) running under Windows. Data were sampled at 10 kHz and low-pass filtered at 2 kHz with a four-pole Bessel filter. The calculated liquid junction potential between intracellular and extracellular solution was also compensated (14.6 mV for normal aCSF, calculated with Patcher’s Power Tools plug-in from http://www.mpibpc.mpg.de/groups/neher/index.php?page=software for IGOR Pro 6 [Wavemetrics, Lake Oswego, OR, USA]).

Whole-cell recordings were performed with pipette solution containing (in mM): 141 K-gluconate, 10 KCl, 10 HEPES, 0.1 EGTA, 2 MgCl_2_, 3 K-ATP, 0.3 Na-ATP and adjusted to pH 7.2 with KOH. For perforated patch clamp recordings and Ca^2+^ imaging, ATP and GTP were omitted from the pipette solution since ATP might distort the measurement of Ca^2+^ handling properties (Matthews et al., 2013) and to prevent uncontrolled permeabilization of the cell membrane during perforated patch clamp recordings (Lindau and Fernandez, 1986). For Ca^2+^ imaging, EGTA was replaced by 100 or 200 μM fura-2 (pentapotassium salt, F1200, Molecular Probes, OR, USA).

Perforated patch recordings were performed using protocols modified from Fan and Palade (Fan and Palade, 1998) and Sarantopoulos (Sarantopoulos et al., 2004). The patch pipette was tip filled with internal solution and backfilled with β-escin (~30 - 60 μg·ml^−1^; E1378; Sigma) or amphotericin B (~200 μg·ml^−1^; A4888, Sigma) containing internal solution to achieve perforated patch recordings. β-escin was prepared as a stock solution (25 mM in internal solution) and stored up to 1 week at −20°C protected from light. Shortly before the experiments, the stock solution was diluted in intracellular saline to a final concentration. Amphotericin B was dissolved in dimethyl sulfoxide (DMSO; final concentration 0.4 – 0.5%; D8418, Sigma) as described previously (Hess et al., 2013; Kyrozis and Reichling, 1995; Rae et al., 1991) and added to the modified pipette solution shortly before use. The amphotericin B solution was freshly prepared every day. Ionophore containing solutions were stored on ice and replaced every 4 hours as needed. After obtaining a GΩ seal, the perforation process was monitored by continually measuring the access resistance (*R*_a_). Experiments were started after *R*_a_ had reached steady-state, and the action potential amplitude was stable (20 – 30 min). A rupture of the membrane and the resulting conversion to the whole-cell configuration was indicated by abrupt changes in *R*_a_ or action potential amplitude. Such experiments were rejected.

### Imaging set up for fluorometric measurements

The imaging setup for fluorometric measurements consisted of an Imago/SensiCam CCD camera with a 640×480 chip (Till Photonics, Gräfelfing, Germany) and a Polychromator IV (Till Photonics) that was coupled *via* an optical fiber into the upright microscope (microscope details are provided above). The camera and the polychromator were controlled by the software Vision (version 4.0, Till Photonics) running under Windows. The excitation light from the polychromator was reflected onto the cells with a dichroic mirror, and emitted fluorescence was detected through an emission filter. The specific filter and mirror settings for the different experiments are given below. Data were acquired as 80×60 frames using 8×8 on-chip binning. Images were recorded in arbitrary units (AU) and stored and analyzed as 12-bit grayscale images. For analyzing the fluorescence dynamics, the mean AU values of regions of interest (ROIs) from the middle of the cell bodies were used. ‘Background subtraction’ for the time series was performed with equivalent time series of neighboring ROIs.

### Rhodamine-Dextran loading

Loading experiments with fluorescent tetramethylrhodamine-dextran (TRD-40kDa MW 40 kDa, D1842, Life Technologies) were performed in the standard whole-cell or the perforated patch configuration. The neurons were loaded with TRD-40kDa *via* the patch pipette (0.08%) and excited at 557 nm (bandpass: 562 nm, 562/40 BrightLine HC, Art-no. F39-562, AHF Analysentechnik AG, Tübingen, Germany; beamsplitter: 593 nm, HC BS 593, Art-no. F38-593, AHF). Emission was detected through a 593 nm long-pass filter (593/LP BrightLine HC, Art-no. F37-594, AHF). TRD-40kDa-fluorescence is reported as *F/F_0_*, where *F* is the measured fluorescence, and *F_0_* is the averaged fluorescence of 5 consecutive images in on-cell mode (before break-in or perforation of the cell membrane). Frames were taken in intervals of 30 s.

### Fluorimetric Ca^2+^ measurements with fura-2

Intracellular Ca^2+^ concentrations were measured with the Ca^2+^ indicator fura-2, a ratiometric dye suitable for absolute Ca^2+^ concentration determination once calibrated (Grynkiewicz et al., 1985; Poenie, 1990). DA SN neurons were loaded with fura-2 (pentapotassium salt, F1200, Molecular Probes) *via* the patch pipette (100 or 200 μM). Fura-2 was excited at 340 nm, 360 nm, or 380 nm (410 nm dichroic mirror; DCLP410, Chroma, Rockingham, VT, USA). Emitted fluorescence was detected through a 440 nm long-pass filter (LP440).

### Calibration and ‘added buffer approach’

The free intracellular Ca^2+^ concentrations were determined as described by (Grynkiewicz et al., 1985), and the added buffer approach to analyze intracellular Ca^2+^ handling properties was performed as previously described (Neher and Augustine, 1992; Paeger et al., 2017; Pippow et al., 2009). For completeness and notation clarification, these methods are described in detail in the *Appendix*. In brief: The added buffer approach was used in combination with whole-cell and perforated patch clamp recordings and optical Ca^2+^ imaging. The added buffer approach is based on a single-compartment model with the rationale that for measurements of intracellular Ca^2+^ concentrations with Ca^2+^ chelator-based indicators, the amplitude and time course of the signals depends on the concentration of the Ca^2+^ indicator (here fura-2). The indicator acts as an exogenous Ca^2+^ buffer and competes with the endogenous Ca^2+^ buffer(s).

The kinetics of cytosolic Ca^2+^ signals strongly rely on the endogenous and exogenous (added) Ca^2+^ buffers of the cell. The amplitude and decay rate of the free intracellular Ca^2+^ change with increasing exogenous buffer concentration: The amplitude of free Ca^2+^ decreases, and the time constant τ_transient_ of the decay increases. If the buffer capacity of the added buffer is known, the time constant of decay (τ_transient_) can be used to estimate, by extrapolation, the Ca^2+^ signal to conditions, with only endogenous buffers present (−κ_B_ = 1 + κ_S_). The model used for this study (Eq. 4) assumes that the decay time constants τ_transient_ are a linear function of the Ca^2+^ binding ratios (κ_B_ and κ_S_). κ_S_ is determined from the negative x-axis intercept of this plot. The slope of the fit is the inverse of the linear extrusion rate (γ). The point of intersection of the linear fit with the y-axis denotes the endogenous decay time constant τ_endo_ (no exogenous Ca^2+^ buffer in the cell). A detailed description of how to estimate the variance of these parameters is provided in the *Appendix*.

### Tools for data analysis and statistics

Data analysis was performed with Igor Pro 6 (Wavemetrics), Spike2 (CED, Cambridge, UK), R (R Development Core Team [2009], http://www.R-project.org) and Graphpad Prism (version 8; Graphpad Software Inc., La Jolla, CA, USA). All calculations for the determination of EGTA purity, its dissociation constant, and the free Ca^2+^ concentrations in the calibration solutions were performed in R. If not stated otherwise, all calculated values are expressed as means ± SEM (standard error of the mean). Box plots were constructed as follows: The ‘+’ signs show the mean, the horizontal line the median of the data. Whiskers indicate the minimum and maximum values of the data. For pairwise comparisons of dependent and independent normal distributions, paired and unpaired t-tests were used, respectively. For pairwise comparisons of independent, not normal distributions, Mann-Whitney U-tests were used. For multiple comparisons, ANOVA was performed; post hoc pairwise comparisons were performed using t-tests as indicated in the figures. For comparison of the resting Ca^2+^ levels ([Ca^2+^]_i_) over time under whole-cell and β-escin perforated-patch clamp conditions, the regression slope, after linear fitting, was also used. Tests were executed using GraphPad Prism 8 (GraphPad Software Inc., La Jolla, CA, USA). A significance level of 0.05 was accepted for all tests. Significance levels were: * p < 0.05, ** p < 0.01, *** p < 0.001. In all figures, n-values are given in brackets. Exact p-values are reported if p > 0.0001. Scaling, contrast enhancement, and *z*-projections of images were performed with ImageJ v2.0.0. The final figures were prepared with Affinity Designer (Ver. 1.6.1, Serif Ltd).

## Acknowledgments

We thank Helmut Wratil for excellent technical assistance. This work was supported by grant SFB 1218/TP B07 from the Deutsche Forschungsgemeinschaft to PK.

## APPENDIX Measurement of free intracellular Ca^2+^ concentration

The free intracellular Ca^2+^ concentrations were determined as in (Grynkiewicz et al., 1985):

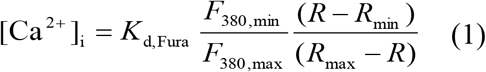

[Ca^2+^]_i_ is the free intracellular Ca^2+^ concentration for the background-subtracted fluorescence ratio *R* from 340 nm and 380 nm excitation. *R*_min_ and *R*_max_ are the ratios at a Ca^2+^ concentration at virtually 0 M (i.e., ideally, no fura-2 molecules are bound to Ca^2+^) and at saturating Ca^2+^ concentrations (i.e., ideally, all fura-2 molecules are saturated with Ca^2+^), respectively. *K*_d,Fura_ is the dissociation constant of fura-2. *F*_380,min◻_/◻*F*_380,max_ is the ratio between the emitted fluorescence of Ca^2+^ free dye and the emitted fluorescence of Ca^2+^ saturated dye at 380 nm excitation, reflecting the dynamic range of the indicator.

The term *K*_d,Fura_ (*F*_380,min_/*F*_380,max_) in Eq. (1) is dependent on the dye concentration and is substituted with the effective dissociation constant *K*_d,Fura,eff_, which is independent of the dye concentration and specific for each experimental setup (Neher, 1989):

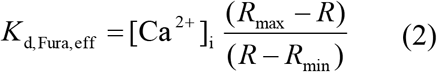

We used *in vitro* calibration (in solution). For calibration *K*_d,Fura,eff_ was determined by measuring fura-2 fluorescence ratios for *R*_max_, *R*_min_, and *R* = *R*_def_. *R*_def_ is the ratio for a defined [Ca^2+^]_i_, which was set to 0.35 μM (see below for the preparation of the solutions). *K*_d,Fura,eff_ was then calculated from Eq. (2). To account for environmental differences between the cytoplasmic milieu and *in vitro* conditions, we used a correction factor (*P*) for *R*_max_ and *R*_min_, as suggested by (Grynkiewicz et al., 1985), see also (Pippow et al., 2009):

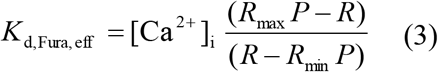

*P* was determined as described by (Poenie, 1990; Zhou and Neher, 1993). First, the fluorescence (peak) deflection at 340 nm was divided by that at 380 nm excitation from voltage-induced intracellular calcium transients (*R*_d,cell_). Second, the ratio (*R*_d,vitro_) from pairs of calibration solutions was determined by dividing (*F*_340,max_ − *F*_340,min_)/(*F*_380,min_ − *F*_380,max_). The correction factor *P* is the fraction of *R*_d,cell_/*R*_d,vitro_.

The fluorescence ratio *R* of an intracellular transient can then be converted to [Ca^2+^]_i_ using:

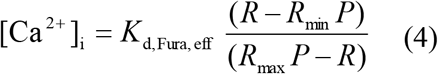

To determine the endogenous Ca^2+^ binding ratio, the dissociation constant (*K*_d,Fura_) has to be known. To formulate a relationship between *K*_d,Fura_ and *K*_d,Fura,eff_ that is independent of the dye concentration, an isocoefficient α was introduced by (Zhou and Neher, 1993):

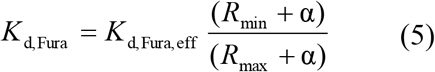

α is the isocoefficient, for which the sum

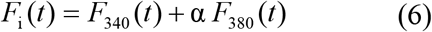

is independent of Ca^2+^ concentration. *F*_340_ and *F*_380_ are the background-subtracted fluorescence signals measured during a brief Ca^2+^ transient. Once the isocoefficient has been determined, *K*_d,Fura_ can be calculated from Eq. (5).

Calibration solutions: The free Ca^2+^ concentrations of the calibration solutions were adjusted by using appropriate proportions of Ca^2+^ and the Ca^2+^ chelator EGTA. The ability of EGTA to bind calcium is highly dependent on the environmental conditions such as ionic strength, temperature, pH, and the concentrations of other metals that compete for binding (Harrison and Bers, 1989, 1987). In theory, the necessary amount of Ca^2+^ and EGTA to set the free Ca^2+^ concentration for the experimental conditions can be computed (Patton et al., 2004). However, small variations in the parameters such as pH, temperature, impurities of chemicals, pipetting, or weighing errors can lead to considerable errors in the estimate of the free Ca^2+^ in EGTA-buffered Ca^2+^ solutions with computer programs (McGuigan et al., 2007). To account for such variations, we determined the free Ca^2+^ concentrations in our calibration solutions by using a Ca^2+^ selective electrode following the guide from McGuigan (McGuigan et al., 1991) and Pippow (Pippow et al., 2009) as described in the supplemental. Calibration solutions were prepared as follows (in mM): *R*_max_: 140 KCl, 2.5 KOH, 15 NaCl, 1 MgCl_2_, 5 HEPES, 10 CaCl_2_ and 0.05 fura-2; *R*_min_: 129.5 KCl, 13 KOH, 15 NaCl, 1 MgCl_2_, 5 HEPES, 8 EGTA and 0.05 fura-2; *R*_def_: 129.5 KCl, 13 KOH, 10.3 NaCl, 4.7 NaOH, 1 MgCl_2_, 5 HEPES, 4 EGTA, 2.7 CaCl_2_ and 0.05 fura-2, yielding a free Ca^2+^ concentration of 0.35 μM. All solutions were adjusted to pH 7.2 with HCl.

The concentrations of free Ca^2+^ were determined from the ratioed imaging signals applying the approach from Grynkiewicz et al. (Grynkiewicz et al., 1985) (Eq. (2)). The calibration was performed *in vitro* (in solution). For calibration, a drop of each calibration solution (200 μl, *R*_min_: no Ca^2+^, *R*_def_: 0.35 μM Ca^2+^, *R*_max_: 10 mM Ca^2+^) was placed on a sylgard coated recording chamber. Ratio images (from 340 nm and 380 nm excitation) were acquired for each solution. The acquired values were: correction factor (Poenie, 1990) *P* = 0.78 ± 0.03, resulting in: *R*_max_ = 1.267 ± 0.055 (*n* = 9); *R*_min_ = 0.116 ± 0.004 (*n* = 9); *R*_def_ = 0.277 ± 0.004 (*n* = 7); *K*_d,Fura,eff_ = 1.048 ± 0.052 μM and isocoefficient α = 0.192 ± 0.003. According to Eq. (5) the *K*_d_ for fura-2 was *K*_d,Fura_ = 0.222 ± 0.082 μM.

## Single compartment model of calcium buffering: the ‘added buffer approach’

For measurements of intracellular Ca^2+^ concentrations with Ca^2+^ chelator-based indicators, the amplitude and time course of the signals are dependent on the concentration of the Ca^2+^ indicator (in this case, fura-2) that acts as an exogenous (added) Ca^2+^ buffer and competes with the endogenous buffers (Helmchen et al., 1997; Helmchen and Tank, 2015; Neher and Augustine, 1992; Tank et al., 1995). With the ‘added buffer approach’ (Neher and Augustine, 1992), the capacity of the endogenous Ca^2+^ buffer in a cell is determined by measuring the decay of the Ca^2+^ signal at different concentrations of ‘added buffer’ and by extrapolating to conditions in which only the endogenous buffer is present. Strictly speaking, the added buffer approach in combination with the whole-cell configuration can only be used to analyze the immobile fraction of Ca^2+^ buffers and only when the measurements start after the mobile Ca^2+^ buffers have diffused out of the cell and do not interfere with the analysis. In contrast, ß-escin perforated recordings can be used to measure the total endogenous Ca^2+^ buffer capacity. The added buffer approach is based on a single-compartment model assuming that a big patch-pipette with a known (clamped) concentration of a Ca^2+^ indicator (fura-2) is connected to a spherical cell. The model assumes that the cell contains an endogenous Ca^2+^ buffer (*S*), of which the total concentration, dissociation constant *K*_d,S_, and its on (*k*^+^) and off (*k*^−^) rates are unknown. When the cell membrane is perforated by β-escin, fura-2 (in its bound and unbound forms) starts to diffuse into the cell, which is considered small enough to be always in a ‘well mixed’ state (i.e., there are no gradients of chemical species within the cell body). In this model three cellular parameters determine the cytosolic Ca^2+^ dynamics: (1) Ca^2+^ sources, (2) Ca^2+^ binding ratio and (3) Ca^2+^ extrusion.

Ca^2+^ sources: In the model, two forms of Ca^2+^ sources are defined: (1) the Ca^2+^ influx from the pipette (*j*_in,pipette_) and (2) the Ca^2+^ entering the cell *via* voltage-activated channels from a brief Ca^2+^ pulse, which can be described by a delta function (*j*_in,stim_) (Helmchen and Tank, 2015; Neher and Augustine, 1992):

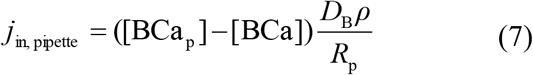

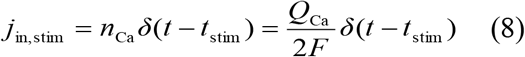

[BCa] is the concentration of the exogenous buffer (fura-2) in its Ca^2+^ bound form (the subscript p is for quantities within the pipette), *D*_B_ is the diffusion coefficient of the exogenous buffer, *ρ* is the specific resistance of the pipette filling solution, *R*_P_ is the pipette resistance, *n*_Ca_ is the Ca^2+^ influx induced by the stimulus, *δ*(*t*−*t*_stim_) is the delta function with *t*_stim_ as the time of stimulus, *Q*_Ca_ is the Ca^2+^ charge influx, and *F* is the Faraday’s constant.

Ca^2+^ binding ratio: In the cell, Ca^2+^ is bound to the Ca^2+^ buffers fura-2 and to the endogenous buffers, which are both assumed to be always in equilibrium with free Ca^2+^ and not saturated. The ability of the experimentally introduced exogenous buffer to bind Ca^2+^ is described by its Ca^2+^ binding ratio that is defined as the ratio of the change in buffer-bound Ca^2+^ over the change in free Ca^2+^:

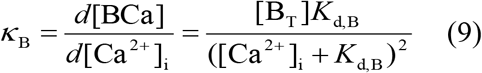

[B_T_] is the total concentration of the exogenous buffer *B* and *K*_d,B_ is its dissociation constant for Ca^2+^. An analogous expression exists for the Ca^2+^ binding ratio of the endogenous buffer *S* (*κ*_S_).

Ca^2+^ Extrusion: the model assumes a linear extrusion mechanism for Ca^2+^ (Neher and Augustine, 1992):

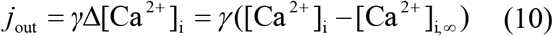

[Ca^2+^]_i,∞_ is the steady-state intracellular Ca^2+^ concentration [Ca^2+^]_i_, to which a transient decays. According to Neher and Augustine (Neher and Augustine, 1992), *γ* ‘reflects the combined action of pumps, exchange carriers, and membrane conductances and has the dimension l s^−1^. In the following, we refer to it as 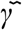. In more recent publications *γ* is defined as the extrusion rate for Ca^2+^ measured in s^−1^ (Helmchen et al., 1997, 1996; Lee et al., 2000a, 2000b; Lips and Keller, 1998; Palecek et al., 1999; Sabatini et al., 2002; Vanselow and Keller, 2000). Accordingly, we use (Jackson and Redman, 2003):

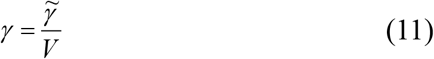

where V is the volume of the cell.

Dynamics of Ca^2+^ transients: the mechanisms described in Eqs. 9 and 10 are combined to the model that describes the dynamics of the Ca^2+^ decay after a brief Ca^2+^ influx:

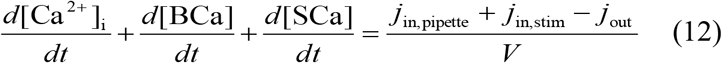

Substituting the time dependent terms for the buffers *B* and *S* with their Ca^2+^ binding ratios *κ*_B_ and *κ*_S_, respectively, [BCa] with its equilibrium value [BCa] = [Ca^2+^]_i_[B_T_]/(*K*_d_+[Ca^2+^]_i_) and 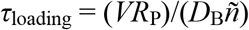 we get (Neher and Augustine, 1992):

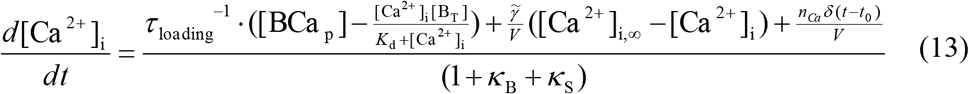

Neher and Augustine (Neher and Augustine, 1992) linearized Eq. (13) to simplify the solution for this equation for the case in which the baseline for [Ca^2+^]_i_ is constant, the concentration of the exogenous buffer [B_T_] is constant and does not influence the baseline Ca^2+^, and when the amplitudes of the Ca^2+^ transients are small (Neher and Augustine, 1992). Where ‘small’ is defined as less than 0.5 *K*_d,Fura_. When the Ca^2+^ pulse is short compared to the decay time constant, the delta function equals zero. Under these conditions, the decay of a voltage-induced Ca^2+^ transient can be described as:

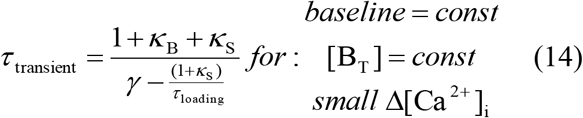

τ_transient_ is proportional to κ_B_, and a linear fit to the data has its negative x-axis intercept at 1 + κ_S_, yielding the endogenous Ca^2+^ binding ratio of the cell. The slope of this fit is the inverse of the Ca^2+^ extrusion rate γ (Eq. (11)). The y-axis intercept yields the time constant τ_endo_ for the decay of the Ca^2+^ transient as it would appear in the cell without exogenous buffer.

## Analysis of calcium buffering

Dye loading was monitored at 360 nm excitation, the isosbestic point of fura-2. Frames were taken at 30 s intervals (3 ms exposure time). We estimated the intracellular fura-2 concentration for different times during the loading curve, assuming that cells were fully loaded when the fluorescence reached a plateau and the fura-2 concentration in the cell and in the pipette is equal (100 or 200 μM) (Lee et al., 2000b).

During fura-2 loading, voltage-activated Ca^2+^ influx was induced by short spike trains, which were elicited by current pulses (500pA, 500ms) from a holding potential of about −70 mV. To monitor the induced Ca^2+^ transients ratiometrically, pairs of images at 340 nm (12 ms exposure time) and 380 nm (4 ms exposure time) excitation were acquired at 10 Hz for ~20 s. Typically 3 – 4 Ca^2+^ transients were induced during loading at different intracellular fura-2 concentrations.

## Ratiometric Data Analysis

## Getting the standard error of the ratiometric estimator

Given measurements at two excitation wavelengths (340 and 380 nm), 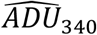 and 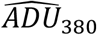 (these are pooled photon counts over the *P* pixels of the ROI) and concomitant measurements 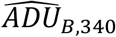 and 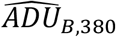 (pooled photon counts over the *P*_*B*_ pixels of the “background region”), what we refer to as the “classical” ratiometric estimator at time *t*_*i*_, is (Joucla et al., 2010) (Eq. 8):

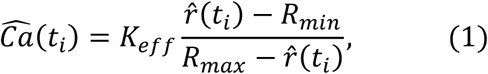

where *K*_*eff*_, *R*_*min*_ and *R*_*max*_ are calibrated parameters (assumed exactly known for now) and where (Joucla et al., 2010) (Eq. 6):

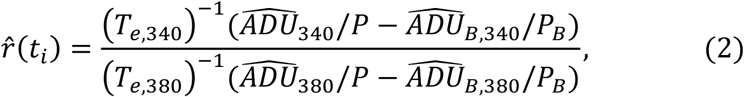

where *T*_*e,λ*_ is the flash duration at wavelength λ. Our model for the fluorescence intensity at each wavelength is (Joucla et al., 2010) (Eq. 2a and 2b):

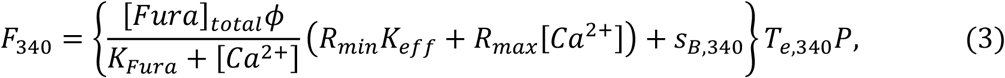

and

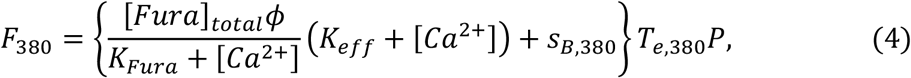

where *S*_*B*,*λ*_ is the auto-fluorescence at wavelength *λ* –assumed homogeneous among the *P* pixels of the ROI–, *K*_*Fura*_ is a calibrated parameter, [*Fura*]_*total*_*ϕ*, is the total (bound plus free) concentration of fura-2 in the cell multiplied by a dimensionless experiment specific parameter, *ϕ*, lumping together the quantum efficiency, the neurite thickness, etc.

Under our assumptions (Joucla et al., 2010) (Eq. 5) we have (we don’t use a “ ⌢ “ here since we are dealing with a *random variable,* not a *realization* of it):

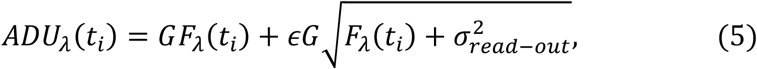

where *G* is the camera gain, 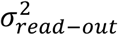 is its read-out variance, *F*_*λ*_(*t*_*i*_) is given by Eq. 3 and 4 and where 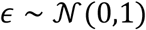 (*ϵ* is a Gaussian random variable with mean 0 and variance 1). In words: *ADU*_*λ*_(*t*_*i*_) has a Gaussian distribution with mean *GF*_*λ*_(*t*_*i*_) and variance 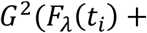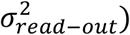

To determine the variance of *ADU*_*λ*_(*t*_*i*_) we need to know *F*_*λ*_(*t*_*i*_) and for that, we need to know [*Ca*^2+^](*t*_*i*_) precisely what we want to estimate. But *EADU*_*λ*_(*t*_*i*_), the expected value of *ADU*_*λ*_(*t*_*i*_), is *GF*_*λ*_(*t*_*i*_) so we can use the observed value 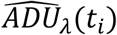 *of ADU*_*λ*_(*t*_*i*_) as an estimate for *GF*_*λ*_(*t*_*i*_) –in Eq. 5 we plug-in 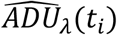 for *GF*_*λ*_(*t*_*i*_) – leading to:

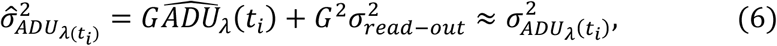

In practice, when we are dealing with several pixels making a ROI, we must multiply the read-out variance by the number of pixels used. With the resulting 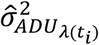 we can work with, to get 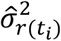 and 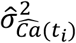. We could use the propagation of uncertainty (or error propagation) together with Eq. 2 and 1 for that, but we can also use a Monte Carlo approach: we draw a thousand quadruple of vectors 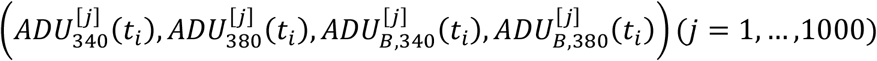 from four independent Gaussian distributions of the general form:

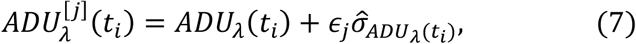

plug-in these quadruples into Eq. 2 giving us 1000 *r*^[*j*]^(*t*_*i*_) before plugging in the latter into Eq. 1 leading to 1000 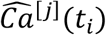. The empirical variance of these simulated observations will be used as 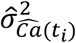.

The method proposed here is slightly less rigorous than the “direct approach” of (Joucla et al., 2010), but it is far more flexible since it does not require an independent estimation or measurement of [*Fura*]_*total*_. In the present study we also chose to consider the calibrated parameters *K*_*eff*_, *R*_*min*_ and *R*_*max*_ as known. Note that the values for these parameters are only estimates (necessarily imprecise since the calibration procedure is subject to errors). However, since the same batch of fura-2 was used for all experiments, they should all have the same (systematic) error. We are not trying to get the exact standard error of the calcium dynamics parameters but to show the difference between two protocols (“classical whole-cell” versus beta-escin perforated whole-cell). Ignoring the uncertainty on the calibrated parameters makes our estimates less variable and helps to make the difference if it exists, clearer. For a true calcium buffering capacity study, the errors on the calibrated parameters should be accounted for, and it would be straightforward to do it with our approach, simply by drawing also *K*_*eff*_, *R*_*min*_ and *R*_*max*_ values from Gaussian distributions centered on their respective estimates with a SD given by the standard errors.

## Objective or “cost” function

We know now how to get both sequences of [*Ca*^2+^] estimates (Eq. 1) and standard errors from sequences of *ADU* measurements. We also consider a calcium dynamics model (Eq. 8-10):

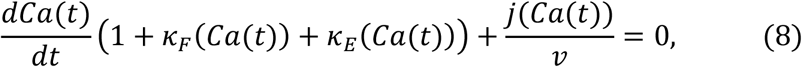

where *Ca*(*t*) stands for [*Ca*^2+^]_*free*_ at time *t*, *v* is the volume of the neurite—within which diffusion effects can be neglected — and

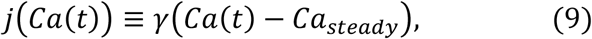

is the model of calcium extrusion — *Ca*_*steady*_ is the steady-state [1] [*Ca*^2+^]_*free*_ — and:

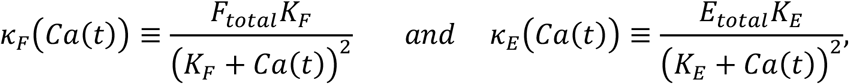

where *F* stands for the fluorophore and *E* for the *endogenous* buffer.

If the values of 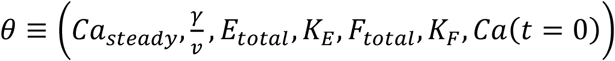 are known, we can substitute these values in Eq. 8, solve it (numerically) and *predict* the time course of calcium at any positive time. Let us write *Ca*(*t*; *θ*) as this solution of Eq. 8. In addition to this model, we have used the results of the previous section, on a set of *N* time points *t*_*i*_, a collection of 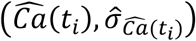. We can then define a first *objective* or *cost* function:

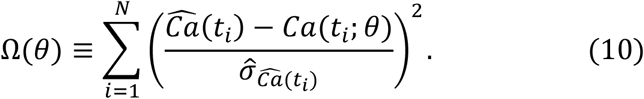

The next task is to find the value of 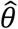 that makes Ω as small as possible; this a classical non-linear weighted least squares problem (see (Nielsen and Madsen, 2010)). The advantage of this setting is that if we can find 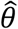 (that is, if the minimum exists and is unique), we also get:

1. Confidence intervals for each element of *θ* through their standard error (in fact, more than that, since we get the full covariance matrix).
2. The distribution from which 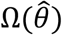 should have been drawn if the model, 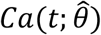, is indeed correct: a *χ*^2^ distribution with *N* − #*θ* degrees of freedom, where #*θ* stands for the number of elements of *θ* that are set from the fitted data.

## Point 2 gives us an objective basis to accept or reject a fit, an essential ingredient for the procedure presented here.

Before going further, we should consider that *θ* in the form just defined contains 7 elements among which, 2 are supposed to be known, *K*_*F*_ and *F*_*total*_. That leaves 5 values to extract from the transient. This is not straightforward since the latter seems, at least at first glance, close to a mono-exponential: 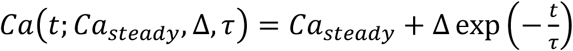 (from which only 3 parameters can be obtained). Therefore, we follow (Neher and Augustine, 1992) and consider an approximation of Eq. 8 where *Ca*(*t*) can be written as *Ca*_*steady*_ + *δCa*(*t*) and *where δCa*(*t*) *is small enough for a first order expansion of Ca*(*t*) *in the vicinity of Ca*_*steady*_ *to be valid*. If we do this first order expansion, neglecting the communication between the pipette and the cytosol during the transient we get (setting *ρ* = 0) in (Neher and Augustine, 1992) (Eq. 15):

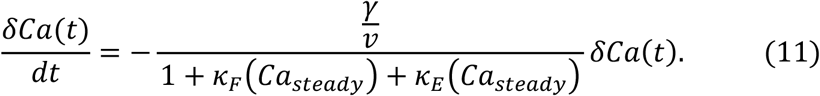

Leading to:

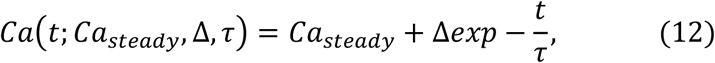

where the time constant *τ* is defined by:

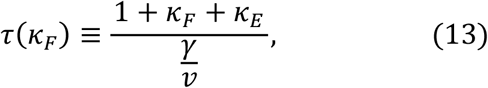

the dependence of the *κ* on *Ca*_*steady*_ has been made implicit. We see that if we can obtain several transients with different values of [*Fura*]_*total*_ leading to several values of *κ*_*F*_, we can exploit the linear dependence of *τ* on both *κ* to estimate *κ*_*E*_ by extrapolation. This is the essence of the added buffer approach. The implementation of the latter requires:

1. Get several (known) values of [*Fura*]_*total*_ and therefore *κ*_*F*_ in the recorded neuron.
2. Trigger a calcium entry at each of these *κ*_*F*_ values and fit *the tail of the transient* with a mono-exponential (Eq. 12)–fitting the tail ensures that the linear approximation of Eq. 8 by Eq. 11 is valid–.
3. Plot *τ* versus *κ*_F_ and fit a straight line to these points. The slope gives 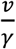 and the x-axis intercept gives *κ*_*E*_.

The fit in point 2 is done with weighted non-linear least squares using *Ca*(*t*; *Ca*_*steady*_, Δ, *τ*) (Eq. 12) instead of *Ca*(*t*; *θ*) in the cost function definition (Eq. 10). Each fit gives 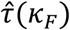 and 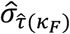 together with a goodness of fit criterion. The 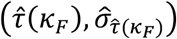 are used in point 3 for a weighted *linear* least squares fit that also leads to confidence intervals on the estimated parameters and to a goodness of fit criterion.

The codes doing the “heavy” part of the analysis were written in C. Some higher-level scripts were written in Python 3. The codes, their documentation, the data, and the way the codes were applied to the data can be found on GitHub (https://github.com/christophe-pouzat/hess-et-al-beta-escin-2019) as well as on zenodo (https://zenodo.org/record/3238248). The weighted linear and non-linear least squares routines, as well as the random number generators, are the ones of the Gnu Scientific Library (http://www.gnu.org/software/gsl/). The http://www.gnu.org/software/gsl/doc/html/nls.html#levenberg-marquardt implementation of the GSL was used for the non-linear least squares. Optimization was ended based on convergence criteria looking at the relative change of each estimated parameter value and at the gradient norm (see the http://www.gnu.org/software/gsl/doc/html/nls.html#testing-for-convergence); as a safeguard, a maximum number of iterations was set, but all the optimizations did actually stop because the convergence criteria were met.

Following Neher and Augustine (1992), our analysis of the calcium transients relies on the linearization (https://en.wikipedia.org/wiki/Linearization) of an assumed calcium dynamics (Eq. 9 to 15, (Neher and Augustine, 1992)). The key hypothesis that is made in all linearizations is that the dynamical system under investigation (here the calcium concentration) is submitted to a small perturbation from its resting state. What constitutes a “small enough” perturbation for the linearization to be valid in practice depends, in the specific case considered here, on parameters we do not know (total endogenous buffer concentration, endogenous buffer dissociation constant, etc.), but we can be sure that the closer we get to baseline, that is the later we look during the transient, the more likely the “small enough” requirement is to be met. If the linearization is valid, the transient (more precisely the portion of the transient that is fitted) must exhibit a mono-exponential decay, something we can rigorously test. In order to get a procedure usable on every transient and considering the fact that the peak response of the first transient can be large - 6 times the baseline value, not really a “small” perturbation - we decided, before seeing the data, to start all the fits from the time at which the transient had lost 50% of its peak value. This is obviously arbitrary but it does not affect our results as the reader can check by downloading our data and our code before rerunning the whole analysis using a different location from which to start the fit. Accordingly, the number of data points used for each fit varies a bit from transient to transient. *This choice of fitting from the “half-peak time” was made before seeing the data*. It is important to realize that once such a choice has been made, we are constrained, the fit is done automatically, and the residual sum of squares (RSS) is compatible or not with the theoretical *χ*^2^ distribution.

Since we used a triple wavelength (340, 380 *and* 360) protocol for each point of the transient, we have access to an independent estimate of [*Fura*]_*total*_.We saw, as in (Joucla et al., 2010) (Fig. 5), that the latter is not constant during the transients. When doing the *τ* versus *κ*_*F*_ linear regression we therefore computed systematically three regressions using the minimum, the mean and the maximum observed values of [*Fura*]_*total*_ during the transient. We always report the results of regression leading to the smallest RSS (best fit). As the reader can check from our GitHub repository, none of our conclusions depend on this. We emphasize again that since our method generates standard errors for 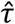, the linear regressions are accepted or rejected on an objective basis (the RSS is compatible or not with the relevant theoretical *χ*^2^ distribution).

---

**Supplement Figure 1.**
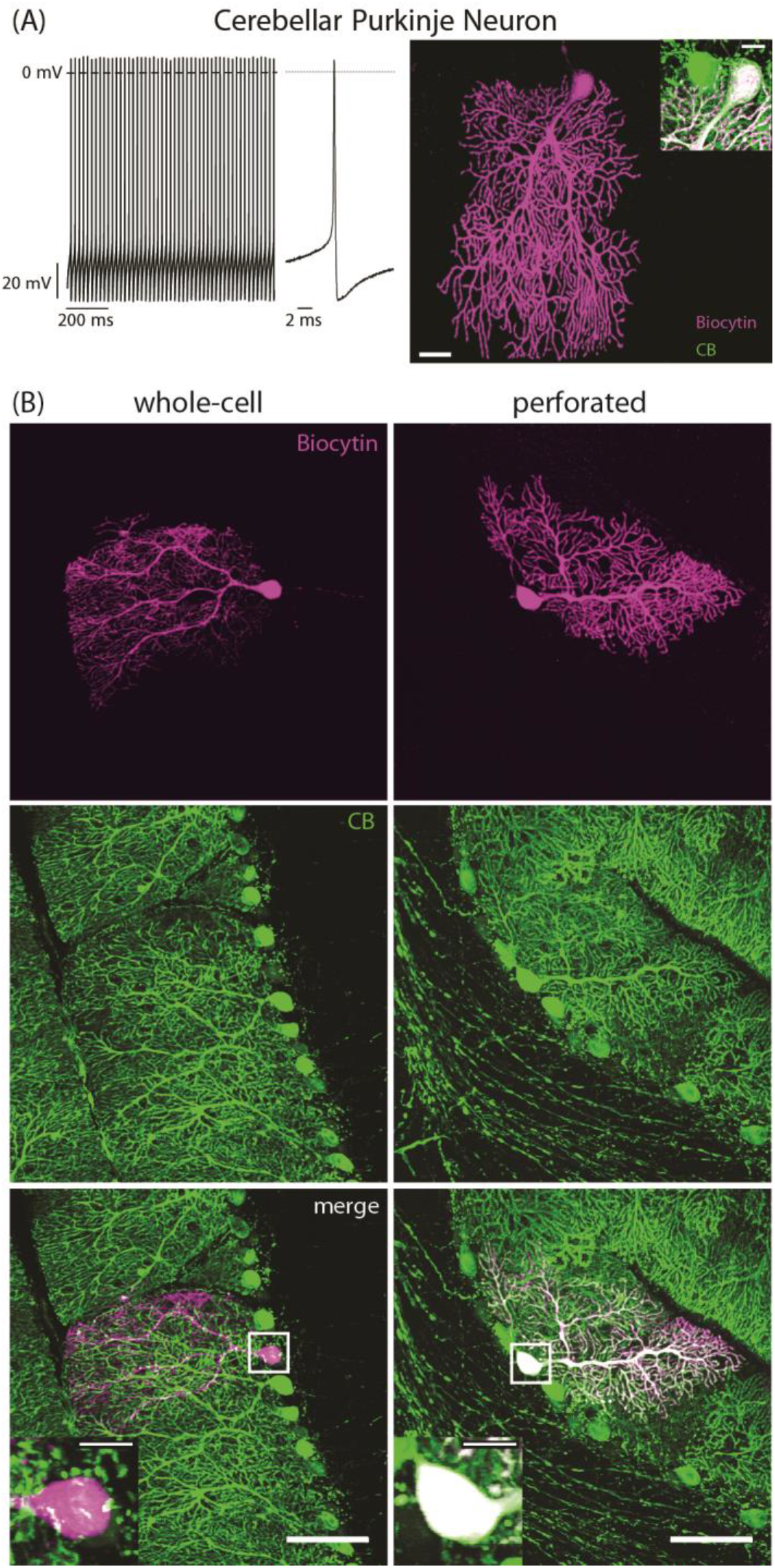
The Ca^2+^ binding protein calbindin D-28k is retained in cerebellar Purkinje neurons during β-escin perforated patch recording. (A) Basic electrophysiological and morphological characteristics of a Purkinje neuron. The neurons were recorded in current clamp and labeled with biocytin-streptavidin (magenta) via the patch pipette. The recording demonstrates the high spontaneous action potential frequency and the short action potential duration, which both are typical for this neuron type. Immunohistochemical labeling demonstrates strong immunoreactivity for calbindin D-28k, which is typical for Purkinje cells (inset). (B) The Ca^2+^ binding protein calbindin D-28k is washed out during whole-cell recording (left column), while it is retained in neurons during β-escin perforated patch recording. (right column). The images show two biocytin-steptavidin labeled Purkinje cells (top panels) and their calbindin D-28k immunoreactivity (middle panels). The overlay of both labels is shown in the bottom panels. Both neurons were recorded for 30 min. Scale bars: 20 μm, insets: 10 μm.

## References

1. Abe H, Watanabe M, Yamakuni T, Kuwano R, Takahashi Y, Kondo H. 1992. Localization of gene expression of calbindin in the brain of adult rats. Neuroscience Letters 138:211–215. doi:10.1016/0304-3940(92)90917-V

2. Akaike N.1994. Glycine responses in rat CNS neurons studied with gramicidin perforated patch recording. The Japanese journal of physiology 44 Suppl 2:S113–8.

3. Akaike N, Harata N. 1994. Nystatin Perforated Patch Recording and Its Applications to Analyses of Intracellular Mechanisms. The Japanese Journal of Physiology 44:433–473. doi:10.2170/jjphysiol.44.433

4. Augustine GJ, Santamaria F, Tanaka K. 2003. Local Calcium Signaling in Neurons. Neuron 40:331–346. doi:10.1016/s0896-6273(03)00639-1

5. Bangham AD, Horne RW, Glauert AM, Dingle JT, Lucy JA. 1962. Action of Saponin on Biological Cell Membranes. Nature 196:196952a0. doi:10.1038/196952a0

6. Bean BP.2007. The action potential in mammalian central neurons. Nature Reviews Neuroscience 8:nrn2148. doi:10.1038/nrn2148

7. Berridge MJ.2011. Calcium Signalling and Alzheimer’s Disease. Neurochem Res 36:1149–1156. doi:10.1007/s11064-010-0371-4

8. Berridge MJ.2006. Calcium microdomains: Organization and function. Cell Calcium 40:405–412. doi:10.1016/j.ceca.2006.09.002

9. Berridge MJ.1998. Neuronal Calcium Signaling. Neuron 21:13–26. doi:10.1016/s0896-6273(00)80510-3

10. Berridge MJ, Lipp P, Bootman MD. 2000. The versatility and universality of calcium signalling. Nature Reviews Molecular Cell Biology 1:11. doi:10.1038/35036035

11. Böttger S, Melzig MF. 2013. The influence of saponins on cell membrane cholesterol. Bioorganic & Medicinal Chemistry 21:7118–7124. doi:10.1016/j.bmc.2013.09.008

12. Chan CS, Gertler TS, Surmeier DJ. 2009. Calcium homeostasis, selective vulnerability and Parkinson’s disease. Trends in Neurosciences 32:249–256. doi:10.1016/j.tins.2009.01.006

13. Ciliax B, Heilman C, Demchyshyn L, Pristupa Z, Ince E, Hersch S, Niznik H, Levey A. 1995. The dopamine transporter: immunochemical characterization and localization in brain. J Neurosci 15:1714–1723. doi:10.1523/JNEUROSCI.15-03-01714.1995

14. Clapham DE.2007. Calcium Signaling. Cell 131:1047–1058. doi:10.1016/j.cell.2007.11.028

1. De Kruijff B, Demel RA. 1974. Polyene antibiotic-sterol interactions in membranes of Acholeplasma laidlawii cells and lecithin liposomes. III. Molecular structure of the polyene antibiotic-cholesterol complexes. Biochimica et Biophysica Acta (BBA) - Biomembranes 339:57–70. doi:10.1016/0005-2736(74)90332-0

16. Delvendahl I, Jablonski L, Baade C, Matveev V, Neher E, Hallermann S. 2015. Reduced endogenous Ca ^2+^ buffering speeds active zone Ca ^2+^ signaling. Proc Natl Acad Sci USA 112:E3075–E3084. doi:10.1073/pnas.1508419112

17. Dodt H-U, Zieglgänsberger W. 1990. Visualizing unstained neurons in living brain slices by infrared DIC-videomicroscopy. Brain Research 537:333–336. doi:10.1016/0006-8993(90)90380-t

18. Fan J-S, Palade P. 1998. Perforated Patch Recording with β-escin. Pflügers Archiv 436:1021–1023. doi:10.1007/pl00008086

19. Fierro L, Llano I. 1996. High endogenous calcium buffering in Purkinje cells from rat cerebellar slices. The Journal of Physiology 496:617–625. doi:10.1113/jphysiol.1996.sp021713

20. Fournet N, Garcia-Segura LM, Norman AW, Orci L. 1986. Selective localization of calcium-binding protein in human brainstem, cerebellum and spinal cord. Brain Research 399:310–316. doi:10.1016/0006-8993(86)91521-0

21. Gilabert JA.2020. Cytoplasmic Calcium Buffering: An Integrative Crosstalk In: Islam MdS, editor. Calcium Signaling. Cham: Springer International Publishing. pp. 163–182. doi:10.1007/978-3-030-12457-1_7

22. Grynkiewicz G, Poenie M, Tsien RY. 1985. A new generation of Ca2+ indicators with greatly improved fluorescence properties. The Journal of biological chemistry 260:3440–50.

23. Harrison SM, Bers DM. 1989. Correction of proton and Ca association constants of EGTA for temperature and ionic strength. American Journal of Physiology-Cell Physiology 256:C1250–C1256. doi:10.1152/ajpcell.1989.256.6.c1250

24. Harrison SM, Bers DM. 1987. The effect of temperature and ionic strength on the apparent Ca-affinity of EGTA and the analogous Ca-chelators BAPTA and dibromo-BAPTA. Biochimica et Biophysica Acta (BBA) - General Subjects 925:133–143. doi:10.1016/0304-4165(87)90102-4

25. Helmchen F, Borst JG, Sakmann B. 1997. Calcium dynamics associated with a single action potential in a CNS presynaptic terminal. Biophysical Journal 72:1458–1471. doi:10.1016/s0006-3495(97)78792-7

26. Helmchen F, Imoto K, Sakmann B. 1996. Ca2+ buffering and action potential-evoked Ca2+ signaling in dendrites of pyramidal neurons. Biophysical Journal 70:1069–1081. doi:10.1016/s0006-3495(96)79653-4

27. Helmchen F, Tank DW. 2015. A Single-Compartment Model of Calcium Dynamics in Nerve Terminals and Dendrites. Cold Spring Harbor Protocols 2015:pdb.top085910. doi:10.1101/pdb.top085910

28. Hess ME, Hess S, Meyer KD, Verhagen LAW, Koch L, Brönneke HS, Dietrich MO, Jordan SD, Saletore Y, Elemento O, Belgardt BF, Franz T, Horvath TL, Rüther U, Jaffrey SR, Kloppenburg P, Brüning JC. 2013. The fat mass and obesity associated gene (Fto) regulates activity of the dopaminergic midbrain circuitry. Nature Neuroscience 16:nn.3449. doi:10.1038/nn.3449

29. Horn R, Marty A. 1988. Muscarinic activation of ionic currents measured by a new whole-cell recording method. The Journal of General Physiology 92:145–159. doi:10.1085/jgp.92.2.145

30. Husch A, Cramer N, Harris-Warrick RM.2011. Long-duration perforated patch recordings from spinal interneurons of adult mice. Journal of neurophysiology 106:2783–9. doi:10.1152/jn.00673.2011

31. Iizuka K, Dobashi K, Yoshii A, Horie T, Suzuki H, Nakazawa T, Mori M. 1997. Receptor-dependent G protein-mediated Ca2+ sensitization in canine airway smooth muscle. Cell Calcium 22:21–30. doi:10.1016/s0143-4160(97)90086-5

32. Iizuka K, Ikebe M, Somlyo AV, Somlyo AP. 1994. Introduction of high molecular weight (IgG) proteins into receptor coupled, permeabilized smooth muscle. Cell Calcium 16:431–445. doi:10.1016/0143-4160(94)90073-6

33. Inoue S, Murata K, Tanaka A, Kakuta E, Tanemura S, Hatakeyama S, Nakamura A, Yamamoto C, Hasebe M, Kosakai K, Yoshino M. 2014. Ionic channel mechanisms mediating the intrinsic excitability of Kenyon cells in the mushroom body of the cricket brain. Journal of Insect Physiology 68:44–57. doi:10.1016/j.jinsphys.2014.06.013

34. Jackson MB, Redman SJ. 2003. Calcium Dynamics, Buffering, and Buffer Saturation in the Boutons of Dentate Granule-Cell Axons in the Hilus. Journal of Neuroscience 23:1612–1621. doi:10.1523/jneurosci.23-05-01612.2003

35. Joucla S, Pippow A, Kloppenburg P, Pouzat C. 2010. Quantitative estimation of calcium dynamics from ratiometric measurements: a direct, nonratioing method. Journal of neurophysiology 103:1130–44. doi:10.1152/jn.00414.2009

36. Klöckener T, Hess S, Belgardt BF, Paeger L, Verhagen LAW, Husch A, Sohn J-W, Hampel B, Dhillon H, Zigman JM, Lowell BB, Williams KW, Elmquist JK, Horvath TL, Kloppenburg P, Brüning JC. 2011. High-fat feeding promotes obesity via insulin receptor/PI3K-dependent inhibition of SF-1 VMH neurons. Nature Neuroscience 14:911. doi:10.1038/nn.2847

37. Kloppenburg P, Zipfel WR, Webb WW, Harris-Warrick RM.2000. Highly Localized Ca2+ Accumulation Revealed by Multiphoton Microscopy in an Identified Motoneuron and Its Modulation by Dopamine. Journal of Neuroscience 20:2523–2533. doi:10.1523/jneurosci.20-07-02523.2000

38. Kobayashi S, Kitazawa T, Somlyo AV, Somlyo AP. 1989. Cytosolic heparin inhibits muscarinic and alpha-adrenergic Ca2+ release in smooth muscle. Physiological role of inositol 1,4,5-trisphosphate in pharmacomechanical coupling. J Biol Chem 264:17997–18004.

39. Konishi M, Watanabe M. 1995. Resting cytoplasmic free Ca2+ concentration in frog skeletal muscle measured with fura-2 conjugated to high molecular weight dextran. The Journal of General Physiology 106:1123–1150. doi:10.1085/jgp.106.6.1123

40. Könner AC, Hess S, Tovar S, Mesaros A, Sánchez-Lasheras C, Evers N, Verhagen LAW, Brönneke HS, Kleinridders A, Hampel B, Kloppenburg P, Brüning JC. 2011. Role for Insulin Signaling in Catecholaminergic Neurons in Control of Energy Homeostasis. Cell Metabolism 13:720–728. doi:10.1016/j.cmet.2011.03.021

41. Korn SJ, Horn R. 1989. Influence of sodium-calcium exchange on calcium current rundown and the duration of calcium-dependent chloride currents in pituitary cells, studied with whole cell and perforated patch recording. The Journal of General Physiology 94:789–812. doi:10.1085/jgp.94.5.789

42. Kruijff BD, Gerritsen WJ, Oerlemans A, Demel RA, Deenen LLM van. 1974. Polyene antibiotic-sterol interactions in membranes of Acholeplasma laidlawii cells and lecithin liposomes. I. Specificity of the membrane permeability changes induced by the polyene antibiotics. Biochimica et Biophysica Acta (BBA) - Biomembranes 339:30–43. doi:10.1016/0005-2736(74)90330-7

43. Kuchibhotla KV, Goldman ST, Lattarulo CR, Wu H-Y, Hyman BT, Bacskai BJ. 2008. Aβ Plaques Lead to Aberrant Regulation of Calcium Homeostasis In Vivo Resulting in Structural and Functional Disruption of Neuronal Networks. Neuron 59:214–225. doi:10.1016/j.neuron.2008.06.008

44. Kyrozis A, Reichling DB. 1995. Perforated-patch recording with gramicidin avoids artifactual changes in intracellular chloride concentration. Journal of Neuroscience Methods 57:27–35. doi:10.1016/0165-0270(94)00116-x

45. Lacey M, Mercuri N, North R. 1989. Two cell types in rat substantia nigra zona compacta distinguished by membrane properties and the actions of dopamine and opioids. Journal of Neuroscience 9:1233–1241. doi:10.1523/jneurosci.09-04-01233.1989

46. Lee S, Rosenmund C, Schwaller B, Neher E. 2000a. Differences in Ca2+ buffering properties between excitatory and inhibitory hippocampal neurons from the rat. The Journal of Physiology 525:405–418. doi:10.1111/j.1469-7793.2000.t01-3-00405.x

47. Lee S, Schwaller B, Neher E. 2000b. Kinetics of Ca2+ binding to parvalbumin in bovine chromaffin cells: implications for [Ca2+] transients of neuronal dendrites. The Journal of Physiology 525:419–432. doi:10.1111/j.1469-7793.2000.t01-2-00419.x

48. Lin K-H, Taschenberger H, Neher E. 2017. Dynamics of volume-averaged intracellular Ca ^2+^ in a rat CNS nerve terminal during single and repetitive voltage-clamp depolarizations: Presynaptic [Ca ^2+^] _i_ dynamics at the rat calyx of Held terminal. J Physiol 595:3219–3236. doi:10.1113/JP272773

49. Lindau M, Fernandez JM. 1986. IgE-mediated degranulation of mast cells does not require opening of ion channels. Nature 319:319150a0. doi:10.1038/319150a0

50. Lips MB, Keller BU. 1998. Endogenous calcium buffering in motoneurones of the nucleus hypoglossus from mouse. The Journal of Physiology 511:105–117. doi:10.1111/j.1469-7793.1998.105bi.x

51. Liu X, Shao R, Li M, Yang G. 2014. Edaravone Protects Neurons in the Rat Substantia Nigra Against 6-Hydroxydopamine-Induced Oxidative Stress Damage. Cell Biochemistry and Biophysics 70:1247–1254. doi:10.1007/s12013-014-0048-8

52. Maravall M, Mainen ZF, Sabatini BL, Svoboda K. 2000. Estimating Intracellular Calcium Concentrations and Buffering without Wavelength Ratioing. Biophysical Journal 78:2655–2667. doi:10.1016/s0006-3495(00)76809-3

53. Matthews EA, Schoch S, Dietrich D. 2013. Tuning Local Calcium Availability: Cell-Type-Specific Immobile Calcium Buffer Capacity in Hippocampal Neurons. The Journal of Neuroscience 33:14431–14445. doi:10.1523/jneurosci.4118-12.2013

54. Mattson MP.2007. Calcium and neurodegeneration. Aging Cell 6:337–350. doi:10.1111/j.1474-9726.2007.00275.x

55. McGuigan JAS, Kay JW, Elder HY, Lüthi D. 2007. Comparison between measured and calculated ionised concentrations in Mg2+ /ATP, Mg2+ /EDTA and Ca2+ /EGTA buffers; influence of changes in temperature, pH and pipetting errors on the ionised concentrations. Magnesium research 20:72–81.

56. McGuigan JAS, Lthi D, Buri A. 1991. Calcium buffer solutions and how to make them: A do it yourself guide. Canadian Journal of Physiology and Pharmacology 69:1733–1749. doi:10.1139/y91-257

57. McKay BE, Turner RW. 2005. Physiological and morphological development of the rat cerebellar Purkinje cell. The Journal of Physiology 567:829–850. doi:10.1113/jphysiol.2005.089383

58. McKay BE, Turner RW. 2004. Kv3 K+ channels enable burst output in rat cerebellar Purkinje cells. European Journal of Neuroscience 20:729–739. doi:10.1111/j.1460-9568.2004.03539.x

59. Müller A, Kukley M, Stausberg P, Beck H, Müller W, Dietrich D. 2005. Endogenous Ca2+ buffer concentration and Ca2+ microdomains in hippocampal neurons. J Neurosci 25:558–565. doi:10.1523/JNEUROSCI.3799-04.2005

60. Murru S, Hess S, Barth E, Almajan ER, Schatton D, Hermans S, Brodesser S, Langer T, Kloppenburg P, Rugarli EI. 2019. Astrocyte◻specific deletion of the mitochondrial m◰AAA protease reveals glial contribution to neurodegeneration. Glia. doi:10.1002/glia.23626

61. Myers VB, Haydon DA. 1972. Ion transfer across lipid membranes in the presence of gramicidin A. Biochimica et Biophysica Acta (BBA) - Biomembranes 274:313–322. doi:10.1016/0005-2736(72)90179-4

62. Neher E, Augustine GJ. 1992. Calcium gradients and buffers in bovine chromaffin cells. The Journal of Physiology 450:273–301. doi:10.1113/jphysiol.1992.sp019127

63. Neuhoff H, Neu A, Liss B, Roeper J. 2002. Ih Channels Contribute to the Different Functional Properties of Identified Dopaminergic Subpopulations in the Midbrain. Journal of Neuroscience 22:1290–1302. doi:10.1523/jneurosci.22-04-01290.2002

64. Nielsen H, Madsen K. 2010. Introduction to Optimization and Data Fitting.

65. Orta G, Ferreira G, José O, Treviño CL, Beltrán C, Darszon A. 2012. Human spermatozoa possess a calcium◻dependent chloride channel that may participate in the acrosomal reaction. The Journal of Physiology 590:2659–2675. doi:10.1113/jphysiol.2011.224485

66. Paeger L, Pippow A, Hess S, Paehler M, Klein AC, Husch A, Pouzat C, Brüning JC, Kloppenburg P. 2017. Energy imbalance alters Ca2+ handling and excitability of POMC neurons. eLife 6:e25641. doi:10.7554/elife.25641

67. Palecek J, Lips MB, Keller BU. 1999. Calcium dynamics and buffering in motoneurones of the mouse spinal cord. The Journal of Physiology 520:485–502. doi:10.1111/j.1469-7793.1999.00485.x

68. Patton C, Thompson S, Epel D. 2004. Some precautions in using chelators to buffer metals in biological solutions. Cell Calcium 35:427–431. doi:10.1016/j.ceca.2003.10.006

69. Pippow A, Husch A, Pouzat C, Kloppenburg P. 2009. Differences of Ca2+ handling properties in identified central olfactory neurons of the antennal lobe. Cell Calcium 46:87–98. doi:10.1016/j.ceca.2009.05.004

70. Poenie M.1990. Alteration of intracellular Fura-2 fluorescence by viscosity: A simple correction. Cell Calcium 11:85–91. doi:10.1016/0143-4160(90)90062-y

71. Rae J, Cooper K, Gates P, Watsky M. 1991. Low access resistance perforated patch recordings using amphotericin B. Journal of Neuroscience Methods 37:15–26. doi:10.1016/0165-0270(91)90017-t

72. Richards CD, Shiroyama T, Kitai ST. 1997. Electrophysiological and immunocytochemical characterization of GABA and dopamine neurons in the substantia nigra of the rat. Neuroscience 80:545–557. doi:10.1016/s0306-4522(97)00093-6

73. Russell JM, Eaton DC, Brodwick MS. 1977. Effects of nystatin on membrane conductance and internal ion activities inAplysia neurons. J Membrain Biol 37:137–156. doi:10.1007/BF01940929

74. Sabatini BL, Oertner TG, Svoboda K. 2002. The Life Cycle of Ca2+ Ions in Dendritic Spines. Neuron 33:439–452. doi:10.1016/s0896-6273(02)00573-1

75. Sarantopoulos C, McCallum JB, Kwok W-M, Hogan Q. 2004. β-escin diminishes voltage-gated calcium current rundown in perforated patch-clamp recordings from rat primary afferent neurons. Journal of Neuroscience Methods 139:61–68. doi:10.1016/j.jneumeth.2004.04.015

76. Sirtori CR.2001. Aescin: pharmacology, pharmacokinetics and therapeutic profile. Pharmacological Research 44:183–193. doi:10.1006/phrs.2001.0847

77. Smits SM, Oerthel L von, Hoekstra EJ, Burbach JPH, Smidt MP. 2013. Molecular Marker Differences Relate to Developmental Position and Subsets of Mesodiencephalic Dopaminergic Neurons. PLoS ONE 8:e76037. doi:10.1371/journal.pone.0076037

78. Tajima Y, Ono K, Akaike N. 1996. Perforated patch-clamp recording in cardiac myocytes using cation-selective ionophore gramicidin. American Journal of Physiology-Cell Physiology 271:C524–C532. doi:10.1152/ajpcell.1996.271.2.C524

79. Tank D, Regehr W, Delaney K. 1995. A quantitative analysis of presynaptic calcium dynamics that contribute to short-term enhancement. Journal of Neuroscience 15:7940–7952. doi:10.1523/jneurosci.15-12-07940.1995

80. Ungless MA, Whistler JL, Malenka RC, Bonci A. 2001. Single cocaine exposure in vivo induces long-term potentiation in dopamine neurons. Nature 411:583. doi:10.1038/35079077

81. Urry DW.1971. The Gramicidin A Transmembrane Channel: A Proposed π(L,D) Helix. Proceedings of the National Academy of Sciences 68:672–676. doi:10.1073/pnas.68.3.672

82. Vanselow BK, Keller BU. 2000. Calcium dynamics and buffering in oculomotor neurones from mouse that are particularly resistant during amyotrophic lateral sclerosis (ALS)◻ related motoneurone disease. The Journal of Physiology 525:433–445. doi:10.1111/j.1469-7793.2000.t01-1-00433.x

83. Ye JH, Zhang J, Xiao C, Kong J-Q. 2006. Patch-clamp studies in the CNS illustrate a simple new method for obtaining viable neurons in rat brain slices: Glycerol replacement of NaCl protects CNS neurons. Journal of Neuroscience Methods 158:251–259. doi:10.1016/j.jneumeth.2006.06.006

84. Zhou Z, Neher E. 1993. Mobile and immobile calcium buffers in bovine adrenal chromaffin cells. The Journal of Physiology 469:245–273. doi:10.1113/jphysiol.1993.sp019813

